# Cholecystokinin input from the anterior cingulate cortex to the lateral periaqueductal gray mediates nocebo pain behavior in mice

**DOI:** 10.1101/2025.02.04.636522

**Authors:** Sandra J. Poulson, Aleksandrina Skvortsova, Fatama T. Zahra, Damien C. Boorman, Seyed A. Karimi, Lisiê V. Paz, Wanning Cui, Antonietta Mandatori, Jacob Burek, Zahra Siddiqi, Maryam Fazili, Shivani R. Gami, Oakley B. Morgan, Melanie Di Maria, Anton Dinh, Lianfang Liang, Robert Contofalsky, Jeffrey S. Mogil, Loren J. Martin

## Abstract

The nocebo effect, the evil twin of the better-known placebo effect, in which anticipation of harm worsens pain and other symptoms, is increasingly thought to be responsible for poor clinical outcomes. In humans, nocebo hyperalgesia (i.e., increased pain sensitivity) is blocked by proglumide, a cholecystokinin (CCK) receptor antagonist. Yet, the neural circuitry underlying nocebo hyperalgesia remains unidentified, largely due to a lack of appropriate animal models. Independently, our two laboratories developed unique animal models of CCK-dependent nocebo hyperalgesia in which the expectation of pain was elicited by environmental or social cues. We find that both nocebo paradigms share a neural circuit involving CCK release from neurons projecting from the anterior cingulate cortex to the lateral periaqueductal gray. This previously unrecognized pathway could represent a promising target for therapeutic interventions in pain-related disorders.

**One-Sentence Summary:** Pain expectations, whether environmentally conditioned or socially transmitted, are mediated by a shared neural circuit involving cholecystokinin (CCK) projections from the anterior cingulate cortex (ACC) to the lateral periaqueductal gray (PAG).

## Introduction

Environmental, including social, cues are pivotal in shaping our perceptions of and responses to pain, influencing both placebo and nocebo effects(Colloca and Benedetti 2009, Vogtle, Barke et al. 2013, Testa, Rossettini et al. 2023). Placebo analgesia arises when the positive expectation of pain relief triggers authentic analgesic outcomes, even without active painkillers—a concept widely explored in human studies(Benedetti, Mayberg et al. 2005, Tracey 2010, Zunhammer, Spisak et al. 2021) and recently in animals(Chen, Goldstein et al. 2024, Chen, Niehaus et al. 2024, Neyama, Wu et al. 2025). In the nocebo effect as it relates to pain, negative expectancies—created either by verbal suggestion, prior experience, or other contextual factors—lead to pain hypersensitivity or hyperalgesia(Colloca 2024). Seminal work on nocebo hyperalgesia in humans by Benedetti and colleagues suggested that it was mediated by the peptide neurotransmitter cholecystokinin (CCK), as the CCK antagonist proglumide was able to abolish the nocebo effect (and potentiate placebo analgesia) in hidden injection experiments(Benedetti, Amanzio et al. 1995, Benedetti, Amanzio et al. 1997). Very little is known about the biology of the nocebo effect beyond this due to the lack of appropriate animal models.

Rodents are capable of acquiring negative pain expectations both through direct associative learning(Martin, Acland et al. 2019, Jee, Zhu et al. 2023) and through social observation(Laviola, Zoratto et al. 2017, Martin, Acland et al. 2019, Smith, Asada et al. 2021), and both forms of learning can produce nocebo hyperalgesia in humans(Vogtle, Barke et al. 2013, Martin, Acland et al. 2019). Here, we describe experiments undertaken independently by two different laboratories, investigating the neurochemistry underlying nocebo hyperalgesia produced either by directly conditioning mice to a context in which they previously received tonic pain(Martin, Acland et al. 2019, Trask, Mogil et al. 2022) or by exposing mice socially to conspecifics that were actively in pain(Laviola, Zoratto et al. 2017). We hypothesized that environmentally conditioned or socially enhanced pain expectations rely on similar neural mechanisms that use CCK as a neurotransmitter. Using a combination of pharmacological, immunohistochemical, viral tracing, and optogenetic approaches, we show that both phenomena depend on CCK receptor activation. Furthermore, we identify a neural circuit involving CCKergic neurons projecting from the anterior cingulate cortex (ACC) to the lateral periaqueductal gray (lPAG) that is crucial for nocebo hyperalgesia across paradigms.

## Results

### Inhibition of CCK receptors blocks nocebo pain hypersensitivity in mice

In humans, conditioning-based negative expectations have been shown to induce nocebo hyperalgesia(Colloca, Petrovic et al. 2010, Brascher, Kleinbohl et al. 2017, Tu, Park et al. 2019). Thus, we posited that prior exposure to pain coupled with subsequent presentation of cues associated with that pain would generate a negative expectation, leading to heightened pain sensitivity in mice, a phenomenon dependent on the CCK system. To test this, we used a conditioned pain hypersensitivity protocol that involved a single pairing of an unconditioned stimulus (UCS; hind paw incision, Fig. 1a) with a conditioned stimulus (CS; environmental context). In our protocol, mice were then returned to the same context 6 days later, after surgical incision sensitivity had resolved (Supplementary Fig. 1), to assess whether re-exposure to the pain-paired context was sufficient to elicit conditioned pain hypersensitivity. When unilateral hind paw incision pain was used as the UCS, robust mechanical hypersensitivity was observed after mice were returned to the same (Fig. 1b), but not different context (Fig. 1c), an effect that was also observed for thermal hypersensitivity (Supplementary Fig. 2). Context-dependent hypersensitivity was blocked by systemic administration of proglumide, a non-specific CCK receptor antagonist (Fig. 1d), and by the CCK-2–selective antagonist LY 225910 (Fig. 1e), consistent with a CCK-dependent contextual nocebo–like effect. Conditioned hypersensitivity was also present in the contralateral hind paw and was abolished by both antagonists (Supplementary Fig. 3), with the effect of LY 225910 replicated in CD-1 mice in an independent experiment (Supplementary Fig. 4).

**Fig. 1.**
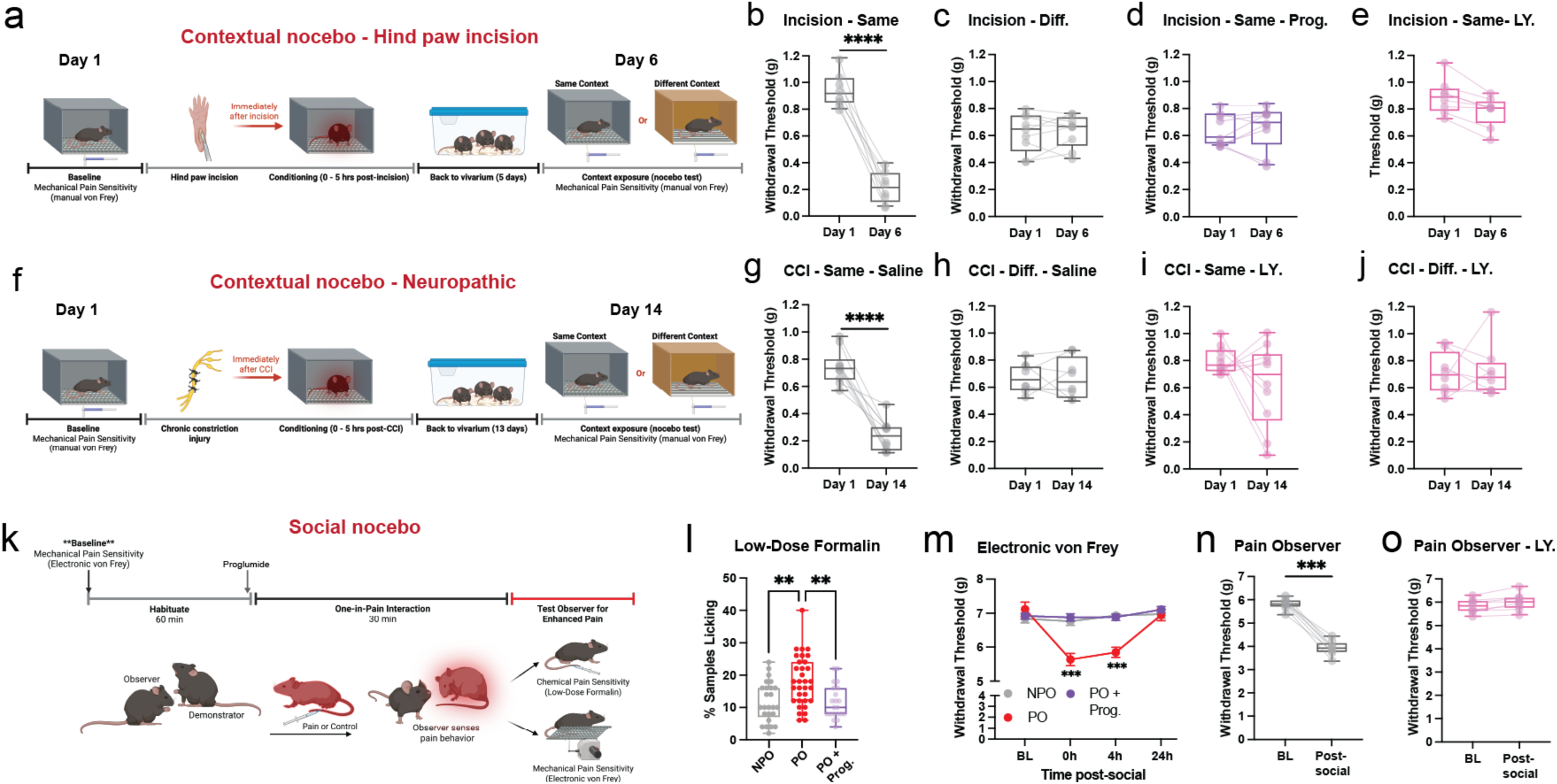
Contextual and social nocebo pain behavior in mice depends on cholecystokinin receptors. **a** Timeline of contextual nocebo conditioning when hind paw incision is used as the unconditioned stimulus (UCS). Mechanical withdrawal thresholds (manual von Frey) for mice test in the same context (**b**, n=9), a different context (**c**, n=9), the same context following proglumide (prog.) treatment (**d**, n=9), or the same context following LY 225910 treatment (**e**, n=8) before and after (Day 6) hind paw incision conditioning. **f** Timeline of contextual nocebo conditioning in a neuropathic pain model using chronic constriction injury (CCI) as the UCS. Mechanical withdrawal thresholds (manual von Frey) for mice test in the same context (**g**, n=10), a different context (h, n=8), the same context following LY 225910 treatment (**I**, n=10), or the same context following LY 225910 treatment (**j**, n=8) before and after (Day 14) CCI conditioning. **k** Timeline of the social nocebo procedure, in which observer mice interact with a demonstrator experiencing pain, followed by testing of the observer. **l** Chemical sensitivity assessed using a low-dose formalin assay in No Pain Observers (NPO, n=25, 15M,10F), Pain Observers (PO, n=33, 23M,10F), and PO + prog. (n= 21, 12M,9F). **m** Time course of mechanical sensitivity assessed by electronic von Frey at baseline (BL), 0, 4, and 24 h following social interaction in NPO (n=11 mice), PO (n=12 mice), and PO + prog. (n=11 mice) groups. A separate experiment measuring mechanical withdrawal thresholds in Pain Observer (PO) mice at baseline and following social interaction (Post-social) in vehicle-treated controls (n = 8) and mice treated with LY225910 (n = 8). Box plots (**b–e and g–j**) show the median and interquartile range, with dots and connecting lines representing paired measurements from individual mice. Symbols in panel **l** represent mean ± SEM percent positive licking samples for the low-dose formalin test, while symbols in **m** represent mean ± SEM for mechanical withdrawal thresholds. Statistical tests include two-way ANOVA for hind paw incision (panels **b–e**) and low-dose formalin (panel **l)**; two-way ANOVA (condition × day) for mechanical sensitivity (panel **m**) and panels **n–o** (drug × time); three-way ANOVA (context × drug × day) for CCI conditioning (panels **g–j**). \*\**p* < 0.01, \*\*\**p* < 0.001 from Šidák-corrected post hoc comparisons or paired *t*-tests with Bonferroni correction. Statistical details are provided in Source Data and file S1.

This hypersensitivity extinguished by Day 9 with repeated testing, while delaying context retrieval to Day 9 still revealed hypersensitivity which also extinguished with repeated testing sessions (Supplementary Fig. 5). To determine whether extinction could be induced by context re-exposure alone, separate cohorts were returned to the pain-paired context without test stimulus application prior to assessment on Day 6 (Supplementary Fig. 6a). Prolonged re-exposure (5 h per day) abolished context-dependent hypersensitivity, whereas shorter re-exposure (3 h per day) produced only a partial attenuation of the response (Supplementary Fig. 6b,c).

We next asked whether chronic constriction injury (CCI) of the sciatic nerve, a unilateral peripheral nerve injury producing neuropathic pain, could similarly produce conditioned pain hypersensitivity when mice were returned to the same context 14 days after injury (Fig. 1f). As with hind paw incision, mice tested in the CCI-paired context showed a significant reduction in mechanical withdrawal thresholds at Day 14 (Fig. 1g), whereas no effect was observed in mice placed in a different context (Fig. 1h). Context-dependent hypersensitivity following CCI conditioning was likewise prevented by administration of the CCK-2–selective antagonist LY 225910 (Fig. 1i,j), and, as in the incision model, extended to the contralateral paw (Supplementary Fig. S7). Finally, independent cohorts showed no sex differences after either hind paw incision (Supplementary Fig. 8a–c) or CCI (Supplementary Fig. 8d–f) conditioning.

Since social cues such as witnessing others in pain also trigger nocebo hyperalgesia in humans (Vogtle, Barke et al. 2013, Saunders, Colagiuri et al. 2023) and facilitate pain in animals (Langford, Crager et al. 2006, Li, Lu et al. 2014, Martin, Hathaway et al. 2015, Lidhar, Darvish-Ghane et al. 2021), we next sought to determine whether the CCK system also mediates the social enhancement of pain in mice. To address this, we adapted a one-in-pain social interaction assay in which a naïve mouse (i.e., Pain Observer, PO) interacted with a cagemate displaying signs of visible pain following an intraplantar injection of formalin (i.e., Pain Demonstrator, PD). PO mice were then subsequently tested for pain responses (either low-dose formalin or mechanical sensitivity) in the absence of PD mice (Fig. 1k). After social interaction, PO mice exhibited increased hind paw licking in response to a low-dose formalin injection compared to No Pain Observer (NPO) mice that interacted with a cagemate injected with saline (i.e., No Pain Demonstrator, NPD). Increased sensitivity in PO mice was blocked by systemic proglumide administration (Fig. 1l). PO mice tested after interacting with a PD showed increased mechanical sensitivity, measured using the electronic von Frey test, immediately and 4 hours after, but not 24 hours after, the social interaction, which was blocked by systemic proglumide (Fig. 1m) or LY 225910 (Fig. 1n,o) given prior to the social interaction. Proglumide administered after the social interaction similarly prevented the increase in pain sensitivity (Supplementary Fig. 9). During the 30-min social interaction, PD mice displayed overt hind paw licking, while NPD mice displayed little to no hind paw licking (Supplementary Fig. 10a). During social interaction, active attending behavior was increased in PO mice, which was not affected by proglumide (Supplementary Fig. 10b). However, PO mice showed increased propinquity (i.e., physical proximity) toward their social partner, which was blocked by proglumide (Supplementary Fig. 10c).

In addition to pain, nocebo is typically accompanied by anxiety, anticipatory fear, and stress(Colloca and Benedetti 2007), raising the possibility that CCK antagonism could reduce hypersensitivity by attenuating a generalized stress-related state. To determine whether CCK signaling contributes more broadly to stress– or fear-related pain modulation, we examined the effects of CCK antagonism in a predator-threat paradigm (Supplementary Fig. 11a) that elicits robust anxiety-like behavior and stress-evoked hypersensitivity(Baumbach, Mui et al. 2025, Baumbach, Mui et al. 2025). In contrast to our nocebo paradigms, CCK antagonism failed to attenuate threat-evoked hypersensitivity (Supplementary Fig. 11b), avoidance (Supplementary Fig. 11c,d), unconditioned freezing to the threat stimulus (Supplementary Fig. 11e) or conditioned freezing upon re-exposure to the threat-paired context (Supplementary Fig. 11f). This suggests that a mild CCK signal is not involved in innate defensive behaviors but is required for expectancy-dependent amplification of pain.

### CCK receptors mediate nocebo hyperalgesia within the lateral periaqueductal gray

The PAG is active during nocebo in human participants, with increased activity observed in the lateral columns of the PAG, more rostral than areas showing placebo-responding activity (Crawford, Mills et al. 2021). For these next experiments, we aimed to demonstrate that activation patterns in the PAG were not specific to any model but rather indicative of the broader nocebo effect. Thus, we used the conditioned pain (i.e., hind paw incision as the UCS) and social pain observation assays to characterize active (i.e., c-Fos^+^) neurons within the PAG (Fig. 2a,b). A survey of the PAG along its rostrocaudal axis (Fig. 2c) revealed similar c-Fos^+^ activation patterns in the contextual and the social nocebo assays. Specifically, the contextual and social nocebo assays showed no change in the precommisural nucleus (Prc), which is the most anterior portion of the PAG (aPAG, Fig. 2d). However, within the intermediate portion (imPAG; bregma –3.6 to – 3.9 mm) of the nucleus, we observed increased c-Fos^+^ neurons in the dorsomedial (dmPAG), dorsolateral (dlPAG) and lateral (lPAG) columns of mice returned to the same context (contextual nocebo) and in PO mice (social nocebo) (Fig. 2e), which was blocked by pre-treatment with proglumide in both paradigms. No apparent difference between the conditions for either assay was observed in the posterior PAG (pPAG; bregma –4.7 to –4.9 mm) in any of the columns (Fig. 2f). Representative images for c-Fos staining are shown for the imPAG for contextual and social nocebo in Fig. 2g,h, respectively. Together, this indicates that the columns of the imPAG may be important for nocebo-induced pain enhancement.

**Fig. 2.**
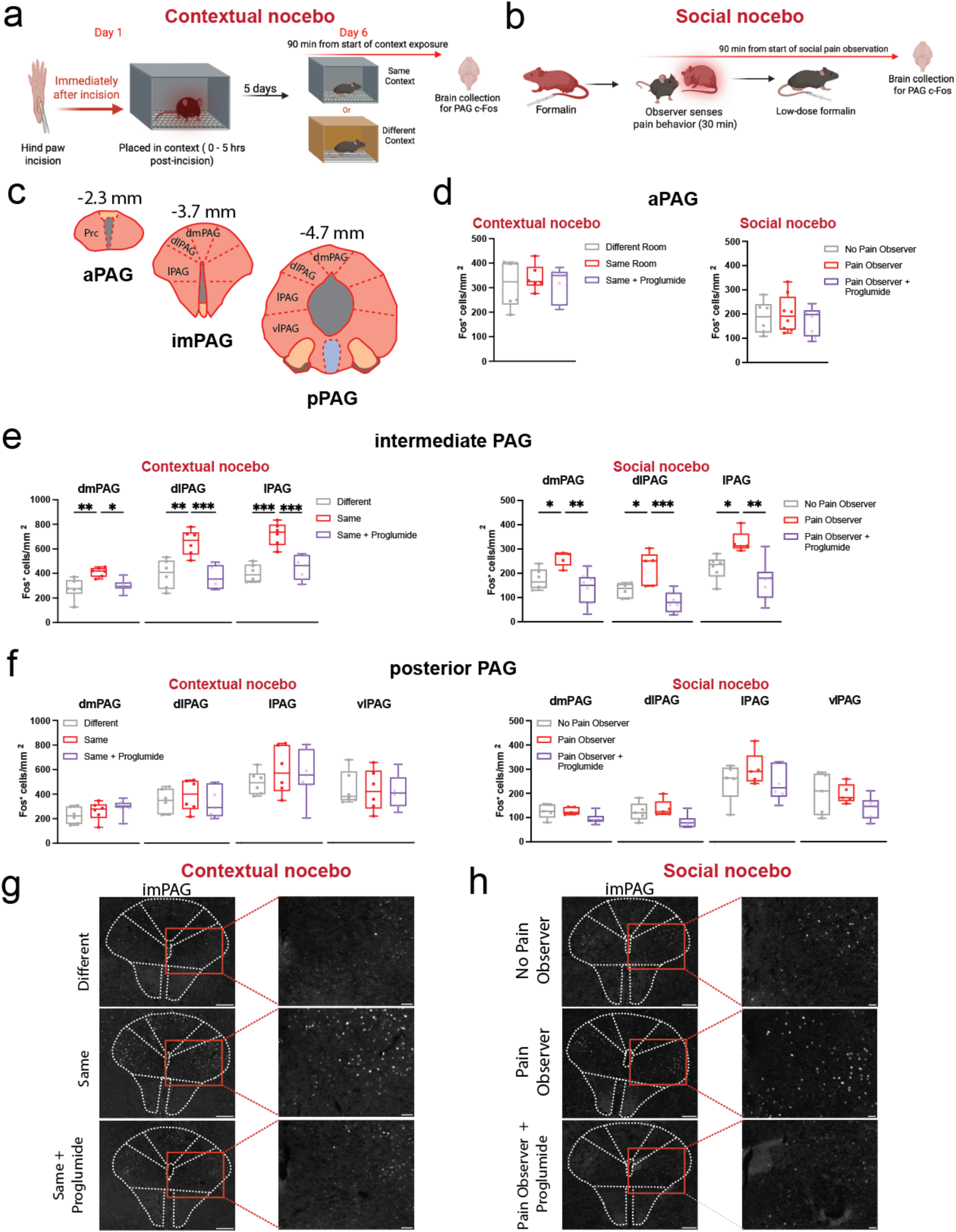
Activation of distinct periaqueductal gray subregions by contextual and social nocebo pain. Timeline of contextual nocebo (**a)** and social nocebo (**b**) pain models used to capture neuronal activation via c-Fos immunohistochemistry. **c** Schematic illustrations of the periaqueductal gray (PAG) showing the bregma levels used for c-Fos quantification, spanning the anterior (aPAG), intermediate (imPAG), and posterior (pPAG) rostrocaudal divisions. **d–f** Quantification of c-Fos-positive cells in the aPAG (**d**), imPAG (**e**), and pPAG (**f**) following contextual and social nocebo pain paradigms. **g** Representative photomicrographs of the imPAG from contextual nocebo groups. **h** Representative photomicrographs of the imPAG for social nocebo groups. Scale bars indicate 200 µm (10x) and 50 µm (zoom) in g, and 160 µm (4x) and 30 µm (zoom) in **h**. For contextual nocebo experiments, n = 6–7 mice/group; for social nocebo experiments, n = 5–8 mice/group. Box plots show the median and interquartile range, with dots representing average c-Fos counts per mm^2^ across three sections per mouse per bregma coordinate from individual mice. Columns of PAG quantification include dorsomedial, dorsolateral, lateral PAG, and ventrolateral (dmPAG, dlPAG, lPAG, and vlPAG, respectively). \**p* < 0.05, \*\**p* < 0.01, \*\*\**p* < 0.001 from Šidák-corrected post hoc comparisons. Statistical details are provided in Source Data and file S1.

Next, we sought to determine whether CCK receptors in the intermediate dmPAG or lPAG were critical for nocebo pain behavior in mice. To do this, we cannulated mice in both regions and tested whether application of proglumide interfered with contextual or social nocebo pain. We also infused CCK-8S into these regions to assess whether activation of CCK receptors was sufficient to evoke pain behavior. The intermediate dmPAG was selected due to its well-established role in sustained fear and anxiety states (Bandler and Depaulis 1991, Borelli and Brandao 2008), while the lPAG was chosen for its involvement in social behavior (Liu, Deng et al. 2022) and pain processing (Kaneko, Oura et al. 2024)—both of which are highly relevant to nocebo pain behavior. Although we observed increased c-Fos^+^ cells in the dlPAG of our nocebo groups, this region is largely associated with active defensive behaviors(Johnson, Lingg et al. 2022), such as escape, which are less relevant to nociceptive processing of social stimuli. Mice infused with proglumide into the intermediate dmPAG continued to show enhanced pain behavior in the contextual and social nocebo models (Fig. 3a–c), while CCK-8S infusion in the intermediate dmPAG did not increase sensitivity to mechanical stimulation (Fig. 3d) or responses on the hot-plate test (Fig. 3e). In contrast, proglumide infusion into the intermediate lPAG blocked pain hypersensitivity in both models (Fig. 3f–h). Importantly, neither systemic proglumide (Supplementary Fig. 12 a,b) nor intra-lPAG proglumide (Supplementary Fig. 12 c) altered baseline nociceptive responses or acute injury-evoked pain in the absence of nocebo conditioning. Application of proglumide to the intermediate lPAG also did not affect distance traveled or velocity in an open field test (Supplementary Fig. 13a,b), indicating that the effects of proglumide on nocebo pain were not simply attributable to altered motor behavior. However, unlike the intermediate dmPAG, CCK-8S microinfusion into the intermediate lPAG increased mechanical sensitivity (Fig. 3i) and pain behavior on the hot-plate test (Fig. 3j) and radiant heat paw-withdrawal test (Supplementary Fig. 14a,b). Differences in drug spread were unlikely to account for our results, as fluorescent muscimol infusions revealed comparable spread within the dmPAG (Supplementary Fig. 15a) and lPAG (Supplementary Fig. 15b) with minimal crossover between PAG columns.

**Figure 3.**
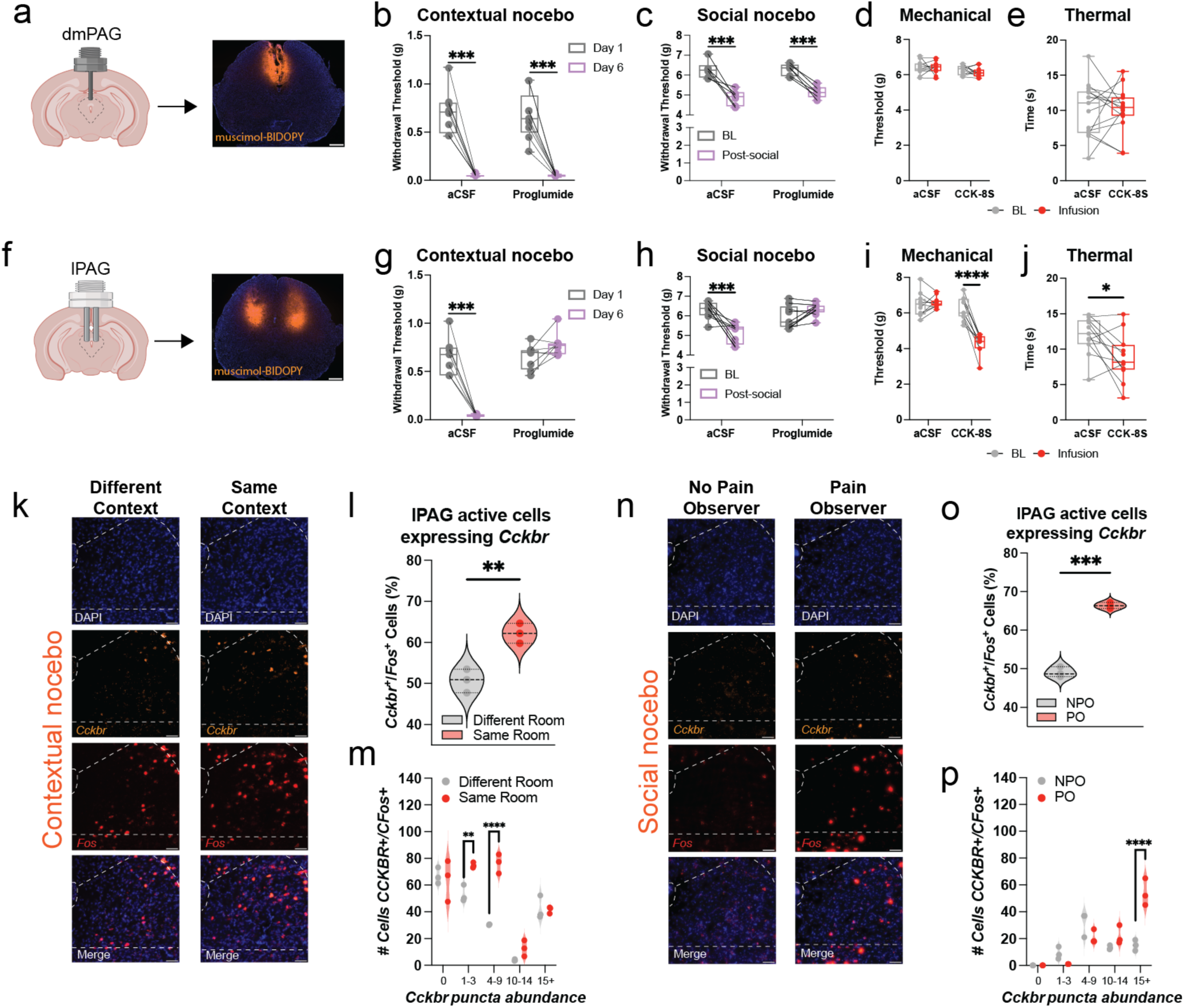
Activation of cholecystokinin receptors in the lateral (but not dorsomedial) periaqueductal gray is necessary for nocebo-induced pain and sufficient to elicit pain-related behavior. **a** Schematic and representative micrograph illustrating cannulation and microinjection targeting the intermediate dorsomedial PAG (dmPAG). For the representative section, fluorescently labeled muscimol (250 nl per side over 5 min) was used to visualize injection spread. Scale bar indicates 100 µm. **b** Mechanical withdrawal thresholds (manual von Frey) for contextual nocebo (hind paw incision) before and after re-exposure to the same context following intra-dmPAG infusion of aCSF or proglumide (n = 8 mice/group). **c** Mechanical withdrawal thresholds (**e**lectronic von Frey) for social nocebo before and after pain observation following intra-dmPAG infusion of aCSF or proglumide (n = 6 mice/group). **d** Intra-dmPAG infusion of CCK-8S does not alter mechanical thresholds as assessed by electronic von Frey testing (n = 11). **e** Intra-dmPAG infusion of CCK-8S does not affect thermal responses on the hot plate (n = 15). **f** Schematic and representative micrograph illustrating cannulation and microinjection targeting the intermediate lateral PAG (lPAG). For the representative section, fluorescently labeled muscimol (250 nl per side over 5 min) was used to visualize injection spread. Scale bar indicates 500 µm. **g** Mechanical withdrawal thresholds (manual von Frey) for contextual nocebo (hind paw incision) before and after re-exposure to the same context following intra-lPAG infusion of aCSF (n = 7 mice) or proglumide (n = 8 mice). **h** Mechanical withdrawal thresholds (**e**lectronic von Frey) for social nocebo before and after pain observation following intra-lPAG infusion of aCSF (n = 9) or proglumide (n = 9). **i** Intra-lPAG infusion of CCK-8S increases mechanical thresholds on the electronic von Frey test (n = 11). **j** Intra-lPAG infusion of CCK-8S reduces latency to lick on the hot plate. CCK-8S infusion in the intermediate lPAG increases mechanical sensitivity on the electronic von Frey test (n = 11). **k** Representative RNAscope images showing *Fos* expression in neurons that contain *Cckbr* mRNA within the intermediate lPAG of mice re-exposed to the same or a different context. Scale bar indicates 100 µm. **l** Violin plots showing quantification of *Cckbr^+^*/*Fos*^+^ neurons in the intermediate lPAG of mice re-exposed to the same or a different (n=3 slices/3 mice/group) context. **m** Bar plots show the number of Fos⁺ neurons in the lPAG expressing increasing *Cckbr* transcript levels, binned by puncta abundance (0, 1–3, 4–9, 10–14, >15 per cell), in mice re-exposed to the same or a different context (n = 3 mice/group). **n** Representative RNAscope images showing *Fos* expression in neurons that contain *Cckbr* mRNA within the intermediate lPAG following social nocebo. Scale bar indicates 100 µm. **o** Violin plots showing quantification of *Cckbr^+^*/*Fos*^+^ neurons in the intermediate lPAG of No Pain Observers (NPO; n=3 slices/3 mice) or Pain Observers (PO; n=3 slices/3 mice). **p** Bar plots show the number of Fos⁺ neurons in the lPAG expressing increasing *Cckbr* transcript levels, binned by puncta abundance (0, 1–3, 4–9, 10–14, >15 per cell), in NPO versus PO mice following the social nocebo paradigm (n = 3 mice/group). Box plots show the median and interquartile range, with dots and connecting lines representing paired measurements from individual mice. Separate two-way ANOVAs (drug × time) for the dmPAG and lPAG for contextual (**b** and **g**) and social nocebo (**c** and **h**); two-way mixed ANOVA (drug X infusion) for mechanical sensitivity (**d** and **i**); two-way mixed ANOVA (cond × puncta group) for RNAscope (**m** and **p**); paired t-tests for RNAscope (**l** and **o**) and hot-plate data (**e** and **j**). \*\**p* < 0.01, \*\*\**p* < 0.001 from Šidák-corrected post hoc comparisons or t-tests where appropriate. Statistical details are provided in Source Data and file S1.

Finally, we asked whether neurons activated during nocebo preferentially expressed the CCK-2 receptor. The CCK-2 receptor, encoded by *Cckbr*, is the predominant CCK receptor subtype in the brain(Moran, Robinson et al. 1986) and is expressed at high levels in the PAG(Lein, Hawrylycz et al. 2007). Using RNAscope, we quantified *Cckbr* transcript abundance in Fos-positive neurons within the lPAG following both contextual and social nocebo paradigms. In the contextual nocebo paradigm, re-exposure to the pain-paired environment increased *Fos* expression in the lPAG, with a higher proportion of activated neurons expressing *Cckbr* compared with mice tested in a different context (Fig. 3k,l). Binning *Fos*⁺ neurons by *Cckbr* RNAscope transcript abundance showed that this increase was driven primarily by recruitment of neurons expressing low-to-moderate levels of *Cckbr* transcripts (Fig. 3m). A similar pattern was observed in the social nocebo paradigm where PO mice exhibited increased *Fos* expression in the lPAG with co-localization of *Cckbr* transcripts (Fig. 3n), along with a higher fraction of Fos⁺ neurons expressing *Cckbr* relative to NPO control mice (Fig. 3o). In contrast to the contextual model, transcript binning in the social paradigm indicated that the increase in *Cckbr* colocalization was driven primarily by Fos⁺ neurons in the highest *Cckbr* abundance bin (15+ puncta), rather than by low-to-moderate expressing cells (Fig. 3p).

To further define the molecular identity of *Cckbr*–expressing neurons in the PAG, we analyzed spatial transcriptomic data from the Allen Brain Cell Atlas(Yao, van Velthoven et al. 2023). *Cckbr* expression in the PAG was largely confined to a small subset of glutamatergic neurons, representing ∼4% of the glutamatergic population, with minimal expression in GABAergic cells (Supplementary Fig. 16a). *Cckbr*⁺ glutamatergic neurons showed extensive co-expression with the μ-opioid receptor gene (*Oprm1)* and the cannabinoid receptor 1 gene (*Cnr1)* with ∼50% expressing both genes within the same cell. In contrast, overlap with stress-related markers such as *Crh* and *Crhr1* was rare (Supplementary Fig. 16b). Spatial mapping showed that *Cckbr*⁺ glutamatergic neurons were distributed across dorsal and lateral regions of the PAG, with *Pou4f1*-associated lineages enriched in the lateral PAG at intermediate and posterior levels (Supplementary Fig. 16c). Quantification across the rostral-caudal axis of the PAG confirmed that *Pou4f1*-derived neurons were most abundant at intermediate levels of the PAG (Supplementary Fig. 16d). Further, *Cckbr*⁺ *Pou4f1* neurons showed prominent co-expression of *Oprm1* and *Cnr1*, positioning this population to modulate descending pain control through established opioid– and endocannabinoid-sensitive PAG circuits (Supplementary Fig. 16e).

### Identification of long-range CCK projections that mediate nocebo behavior

To determine whether the source of CCK within the lPAG arises from projection neurons in other brain regions, we infused a Cre-dependent retrograde AAV expressing mNeptune into the intermediate lPAG of CCK-IRES-Cre mice (Fig. 4a–c). Retrogradely labeled neurons were detected in several cortical regions, including the insula, motor cortex, retrosplenial cortex, and parietal cortex (Fig. 4b–d). However, we identified a distinct and densely labeled population of CCK projection neurons in the anterior cingulate cortex (ACC) (Fig. 4b–e). These ACC neurons were largely confined to layer V and constituted the most prominent source of CCK-positive projections to the intermediate lPAG (Fig. 4e). Quantification across regions confirmed that the ACC showed a significantly higher proportion of retrogradely labeled CCK^+^ neurons relative to total NeuN-positive cells compared to all other regions examined (Fig. 4f). Immunocytochemical analysis revealed that retrogradely labeled ACC^CCK^ neurons co-express EMX1 and exhibit pyramidal morphology, consistent with an excitatory phenotype (Supplementary Fig. 17). This is in line with transcriptomic classification of CCK-expressing ACC neurons (Supplementary Fig. 18).

**Figure 4.**
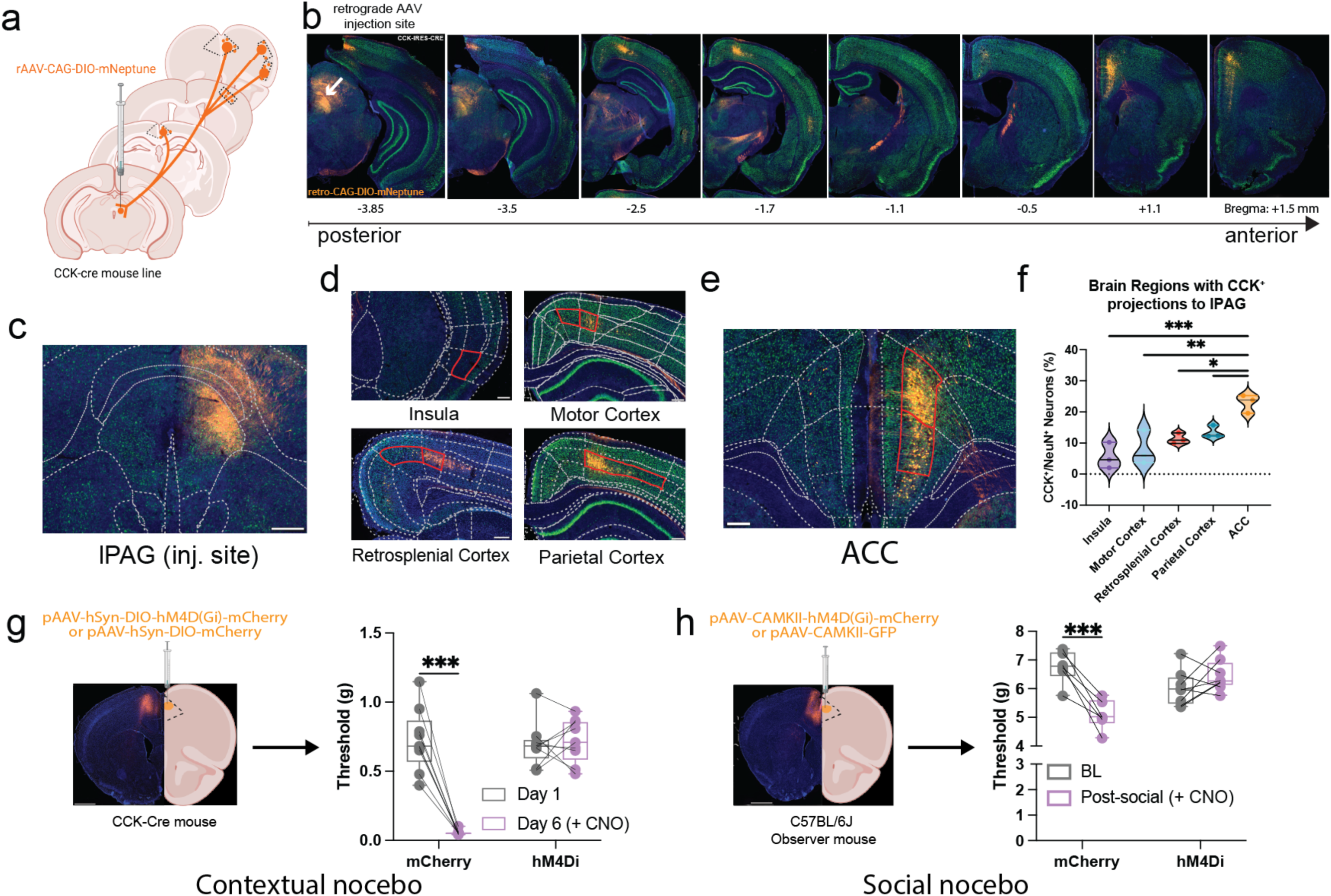
Cholecystokinin-expressing neurons in the anterior cingulate cortex project to the lateral periaqueductal gray and are required for contextual and social nocebo-induced pain hypersensitivity. **a** Schematic illustrating Cre-dependent retrograde viral tracing from the intermediate lateral periaqueductal gray (lPAG) using rAAV-CAG-DIO-mNeptune injected into CCK-Cre mice (n = 3). **b** Representative coronal sections spanning posterior to anterior cortex showing retrogradely labeled CCK+ projection neurons following lPAG injection. Prominent labeling is observed in the anterior cingulate cortex (ACC), with sparser labeling in insular, motor, retrosplenial, parietal, and auditory cortices. Bregma levels are indicated below. **c** Representative micrograph at the lPAG injection site showing co-localization or retrograde mNeptune signal with the neuronal marker NeuN. **d** Higher-magnification micrograph of the ACC showing dense retrograde labeling of CCK⁺ projection neurons. For panels **d** and **e**, NeuN⁺ neurons and NeuN⁺/CCK⁺ neurons were quantified within layer V (red outlines). Scale bars, 200 µm. **f** Violin plots showing the percentage of CCK^+^ neurons relative to the total NeuN^+^ neurons in each cortical region projecting to the lPAG. g *Left:* Experimental schematic for chemogenetic inhibition of CCK⁺ neurons using pAAV-hSyn-DIO-hM4D(Gi)-mCherry or pAAV-hSyn-DIO--mCherry virus in CCK-Cre mice during the contextual nocebo paradigm (n = 9 mice/group). Scale bar, 1 mm. *Right:* Mechanical withdrawal thresholds (manual von Frey) measured on Day 1 and Day 6 following contextual re-exposure, showing that inhibition of CCK⁺ neurons prevents nocebo-induced hypersensitivity (n = 9 mice/group). h Left: Experimental schematic for chemogenetic inhibition of cortical excitatory neurons using pAAV-CAMKII-hM4D(Gi)-mCherry (n = 8 mice) or pAAV-CAMKII-GFP (n = 6 mice) in C57BL/6J observer mice during the social nocebo paradigm. Scale bar,1 mm. *Right:* Mechanical withdrawal thresholds measured at baseline and following social pain observation, demonstrating that inhibition of cortical excitatory neurons attenuates social nocebo-induced hypersensitivity. Box plots show the median and interquartile range, with dots and connecting lines representing paired measurements from individual mice. One-way ANOVA (**f**) or two-way ANOVA (virus × time) for nocebo behavioral tests (g and **h**). \**p*< 0.05, \**p* < 0.01, \*\*\**p* < 0.001 from Šidák-corrected post hoc comparisons. Data are shown as mean ± SEM. Statistical details are provided in Source Data and file S1.

To provide supporting evidence for involvement of the ACC^CCK^→lPAG pathway during nocebo retrieval, we quantified c-Fos expression in retrogradely labeled neurons from a small subset of mice following re-exposure to the same or different context (Supplementary Fig. S19a). Consistent with behavioral reinstatement, context-dependent increases in overall c-Fos expression were observed selectively in the ACC layer 2/3, while enhanced c-Fos co-labeling in retrogradely labeled neurons was observed in ACC layer 5, specifically in the same-context condition (Supplementary Fig. 19b,c). No context-dependent differences in c-Fos or co-labeling with retrogradely labeled neurons were detected in other regions or layers (Supplementary Fig. 19d–f). To directly test whether ACC^CCK^ neurons were required for nocebo behavior, we next used a chemogenetic approach to silence CCK⁺ neurons in the ACC during nocebo retrieval. Inhibition of ACC^CCK^ neurons selectively blocked contextual nocebo responses (Fig. 4g), whereas broader inhibition of excitatory CaMKII-expressing neurons in the ACC disrupted social nocebo responses (Fig. 4h), demonstrating a causal role for ACC^CCK^ and excitatory neurons in mediating nocebo pain behavior.

### ACC^CCK^ → lPAG projections are critical for nocebo pain behavior and pain sensitivity

Based on the microinfusion and viral tracing experiments, we hypothesized that ACC^CCK^ neurons project to the lPAG to mediate nocebo pain behavior in mice. To test this, we targeted ACC^CCK^ terminals in the lPAG using projection-specific optogenetic inhibition. We injected an inhibitory opsin (eOPN3) into the ACC and implanted an optic cannula above the lPAG (Fig. 5a). After 3 weeks of expression, we tested whether optogenetically silencing ACC^CCK^→ lPAG projections disrupted contextual or social nocebo responses. During re-exposure to the pain-paired context, mice expressing a control virus displayed nocebo hypersensitivitiy, whereas optogenetic inhibition of ACC^CCK^→ lPAG terminals blocked contextual nocebo (Fig. 5b). A similar effect was observed in the social nocebo paradigm, where silencing this projection during pain observation prevented the expression of social nocebo (Fig. 5c). We next examined whether ACC^CCK^→lPAG activity was required for contextual nocebo learning. Optogenetic inhibition of this pathway prior to conditioning did not alter baseline mechanical thresholds, and inhibition during the 5-h conditioning period likewise failed to prevent contextual nocebo hypersensitivity when mice were tested without optogenetic inhibition on Day 6 (Fig. 5d). This suggests that suppressing ACC^CCK^→lPAG signaling during baseline or conditioning was not sufficient to disrupt subsequent nocebo expression.

**Figure 5.**
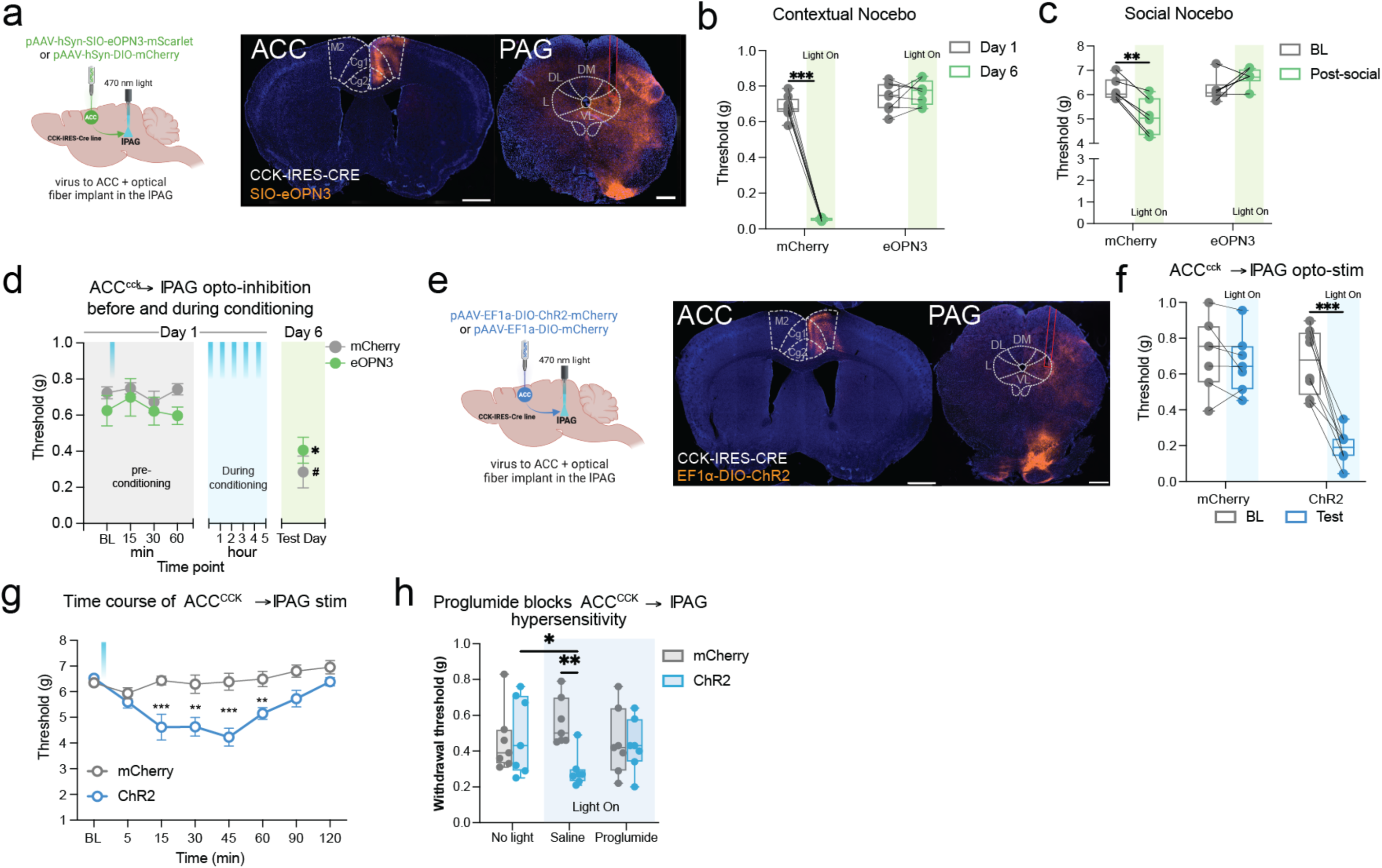
Cholecystokinin-expressing anterior cingulate cortex neurons projecting to the lateral periaqueductal gray are required for nocebo-induced pain hypersensitivity and sufficient to drive pain behavior. **a** Experimental schematic illustrating Cre-dependent expression of the optogenetic inhibitory opsin eOPN3 or control virus (mCherry) in the anterior cingulate cortex (ACC) of CCK-IRES-Cre mice, with optical fiber implantation targeting the lateral periaqueductal gray (lPAG). Mice were tested in contextual or social nocebo paradigms with light delivery to the lPAG. Representative coronal sections show viral expression in the ACC and terminal labeling in the PAG. **b** Mechanical withdrawal thresholds during contextual nocebo (hind paw incision) assessed at baseline (Day 1) and following re-exposure to the conditioned context (Day 6) during optogenetic inhibition of ACC→lPAG projections (Light On) in mCherry and eOPN3 mice (n = 8 mice/group). **c** Mechanical withdrawal thresholds during social nocebo assessed at baseline (BL) and following pain observation (Post-social) during optogenetic inhibition of ACC→lPAG projections (Light On) in mCherry and eOPN3 mice (n = 6 mice/group). **d** Time course of mechanical sensitivity during contextual conditioning, showing that inhibition of ACC→lPAG projections during conditioning does not prevent the development of nocebo-induced hypersensitivity. Shaded regions indicate pre-conditioning, conditioning (Light On), and test phases (n = 8 mCherry mice; n = 6 eOPN3 mice). **e** Experimental schematic illustrating Cre-dependent expression of ChR2 or control virus (mCherry) in the ACC of CCK-IRES-Cre mice with optical fiber implantation targeting the lPAG. Representative coronal sections show viral expression in the ACC and terminal labeling in the PAG. **f** Optogenetic activation of ACC→lPAG projections during testing (Light On) induces mechanical hypersensitivity (manual von Frey) in ChR2-expressing mice(n = 8 mice) but not in mCherry controls (n = 7 mice). **g** Time course of mechanical withdrawal thresholds (electronic von Frey) during sustained optogenetic activation of ACC→lPAG projections, demonstrating rapid onset and time course of hypersensitivity (n = 7 mice/group). **h** Pharmacological blockade of CCK receptors in the lPAG with proglumide attenuates optogenetically evoked hypersensitivity during ACC→lPAG stimulation, indicating that CCK signaling in the lPAG is required for this effect (n = 7 mice/group). Green or blue light shaded regions indicate light on stimulation. Two-way ANOVA (virus X time) for nocebo behavioral tests (**b** and **c**) and ChR2 data (**d**, **f**, **g**, **h**). Box plots show the median and interquartile range, with dots and connecting lines representing paired measurements from individual mice. Data in panels d, g and h are shown as means ± SEM. In panel d, *In panel h, # p<0.05 compared with ChR2 no light baseline, and **p<0.01 compared with mCherry no light baseline. For all other panels \**p* < 0.05, \*\**p* < 0.01, \*\*\**p* < 0.001 from Šidák-corrected or Tukey’s post hoc comparisons. Statistical details are provided in Source Data and file S1.

Because we showed that the ACC^CCK^→lPAG pathway was necessary for contextual and social nocebo, we further aimed to determine whether its activation was sufficient to drive pain-like behavior. To this end, we expressed channelrhodopsin-2 (ChR2) coupled to a mCherry tag in CCK neurons of the ACC and implanted an optical fiber above the lPAG (Fig. 5e), with mCherry-only mice serving as controls. Optical stimulation of ACC^CCK^→lPAG terminals with blue laser light (470 nm, 20 Hz, 10 mW/mm^2^) produced marked mechanical hypersensitivity in ChR2-expressing mice but not controls (Fig 5f). This hypersensitivity emerged rapidly and persisted for approximately 60 min (Fig. 5g), indicating that activation of this projection was sufficient to induce a pain-like state. Finally, administration of the CCK receptor antagonist proglumide attenuated the hypersensitivity evoked by ACC^CCK^→lPAG optogenetic stimulation (Fig. 5h).

## Discussion

Here, we identify a cortico–midbrain circuit in which CCK released from ACC neurons activates CCK-2 receptors in the intermediate lPAG to amplify pain sensitivity under conditions of negative expectation. Across two distinct nocebo paradigms, we find convergent behavioral, pharmacological, anatomical, and circuit-specific evidence that this ACC^CCK^→lPAG pathway is both necessary and sufficient for nocebo-like hyperalgesia in mice. The rigor and generalizability of our findings are enhanced by the independent but converging early work by two different laboratories, unaware of each other’s findings.

In humans, proglumide attenuates nocebo hyperalgesia(Benedetti, Amanzio et al. 1995, Benedetti, Amanzio et al. 1997), and here both non-selective CCK receptor blockade and selective CCK-2 receptor antagonism prevented nocebo hypersensitivity. This effect cannot be explained by a general reduction in nociception or stress responses, as systemic proglumide left baseline sensitivity and formalin-evoked behaviour unchanged, and intra-lPAG proglumide did not affect incision-evoked pain outside of the nocebo context. Likewise, a predator-odor paradigm that produces robust stress-evoked hypersensitivity and avoidance(Baumbach, Mui et al. 2025, Baumbach, Mui et al. 2025) remained intact under proglumide, indicating that a mild CCK signal is not required for general stress-induced hyperalgesia and defensive responses induced by this stimulus. This aligns with human findings that proglumide reduces nocebo hyperalgesia while leaving the associated endocrine stress response unchanged(Benedetti, Amanzio et al. 2006), demonstrating that CCK blockade uncouples pain amplification from stress physiology rather than directly suppressing the stress response. This contrasts with conditions of intense or sustained stress, where social defeat produces widespread increases in brain CCK and engages endocrine responses, and chronic treatment with the selective CCK-2 antagonist CI-988 reduces corticosterone elevation(Becker, Zeau et al. 2008) and stress-enhanced formalin pain(Andre, Zeau et al. 2005). Such effects likely reflect large-scale recruitment of CCK during sustained threat, whereas our paradigms depend on CCK signaling within a defined cortical–PAG circuit. This suggests that CCK recruitment depends on the nature of the threat signal, supporting a model in which CCK is not universally engaged by pain or stress, but can be selectively recruited to amplify nociceptive processing under conditions of negative expectation.

Our unique dataset provides an analysis of the PAG across coronal planes and along the anterior-posterior axis, revealing distinct PAG columns activated by our nocebo paradigms, extending beyond the typically studied posterior PAG. Both, the contextual and social nocebo paradigms increased c-Fos expression in the intermediate lPAG, an effect blocked by proglumide, identifying this region as a crucial hub for CCK-dependent nocebo pain responses. This mirrors findings from human fMRI studies, where neural activation in nocebo responders is localized to the lateral columns of the PAG, more rostral than in placebo responders (Crawford, Mills et al. 2021). Although evidence from animal models in support of the role of the PAG in nocebo hyperalgesia has been limited, our findings, coupled with recent studies of conditioned and socially enhanced nausea behaviors in rats (Zhang, Huang et al. 2024), highlight the significance of the PAG and its interconnected regions, such as the ACC, in nocebo effects. At the cellular level, the contextual nocebo paradigm recruited a broader population of Fos-positive lPAG neurons expressing predominantly low-to-moderate levels of *Cckbr* transcripts, whereas the social nocebo condition engaged fewer neurons but was enriched for cells with high *Cckbr* expression, indicating recruitment of distinct gain states within the same circuit rather than uniform scaling of activity. At intermediate rostrocaudal levels of the PAG, *Cckbr*⁺ neurons were enriched within a population marked by expression of *Pou4f1*, a transcription factor associated with defined neuronal lineages in sensory systems(Zou, Li et al. 2012, Ray, Torck et al. 2018). Although *Pou4f1* has not been directly linked to descending pain modulation, its enrichment within specific *Cckbr^+^* PAG neurons, together with co-expression of *Oprm1* and *Cnr1*, is consistent with engagement of circuits known to regulate nociceptive processing.

Previous work has established that glutamatergic ACC projections to the dlPAG and lPAG contribute to reflexive and active avoidance of pain (Lee, You et al. 2022), but it remains unclear whether these effects reflect broad engagement of ACC output or selective recruitment of defined neuronal populations under specific cognitive conditions. Our findings suggest that nocebo expression depends on selective engagement of a predominantly excitatory, CCK-expressing population of ACC neurons projecting to the intermediate lPAG. Although CCK is expressed in both excitatory and inhibitory cortical neurons(Asim, Wang et al. 2024), several observations indicate that the nocebo effect is largely mediated by an excitatory population. ACC^CCK^→lPAG-projecting neurons were confined to layer V and displayed pyramidal morphology characteristic of corticofugal glutamatergic cells. Retrograde tracing combined with Fos mapping showed selective activation of these projection neurons during nocebo expression rather than broad activation of local cortical networks, and spatial transcriptomic classification further demonstrated that ACC^CCK^ expression is strongly enriched in intratelencephalic (IT)– and extratelencephalic (ET)-type excitatory subclasses, with interneurons representing only a minor fraction. Consistent with this, chemogenetic inhibition of ACC^CCK^ or ACC^CaMKII^ neurons, or optogenetic inactivation of ACC^CCK^ terminals in the lPAG blocked nocebo hypersensitivity. Conversely, optogenetic activation was sufficient to induce mechanical allodynia, which was blocked by proglumide. This may indicate that CCK release from ACC terminals acts as a gain-modulating signal that biases nociceptive output through enhanced glutamate release(Deng, Xiao et al. 2010), consistent with anti-opioid and pain-facilitatory roles of CCK signaling(Hebb, Poulin et al. 2005, Benedetti, Amanzio et al. 2011, Bernard, Danigo et al. 2021). Inhibition of the ACC^CCK^→lPAG pathway during conditioning did not prevent subsequent nocebo expression, indicating that this pathway is recruited during expression rather than acquisition of contextual nocebo associations.

Placebo and nocebo effects are often viewed as independent because they engage different neural and biochemical pathways. Placebo analgesia, for instance, can be disrupted by the opioid receptor antagonist, naloxone, demonstrating the involvement of endogenous opioids(Levine, Gordon et al. 1978). Conversely, in human models of nocebo hyperalgesia, increased pain responses are not reversed by naloxone(Benedetti, Amanzio et al. 1997). Interestingly, application of the CCK-2 receptor agonist, pentagastrin, disrupts placebo analgesia in humans(Benedetti, Amanzio et al. 2011), whereas proglumide enhances placebo responses(Benedetti, Amanzio et al. 1995). Recent circuit-level studies of placebo analgesia indicate that expectation engages a descending inhibitory pathway in which cortical signals recruit ventrolateral PAG neurons that project to the rostroventromedial medulla and suppress nociceptive transmission through endogenous opioid signaling(Kimmey, Ejoh et al. 2025, Livrizzi, Liao et al. 2025, Neyama, Wu et al. 2025). In these models, cortical input from medial prefrontal cortex (mPFC) and ACC gates activity of vlPAG projection neurons, producing analgesia only when expectations of pain-relief or lower pain are present rather than through non-specific activation of the PAG. Positive expectation therefore preferentially engages opioid-dependent vlPAG output that suppresses nociceptive transmission, while negative expectation recruits a CCK-sensitive population in the intermediate lPAG that facilitates pain processing. Rather than activating separate modulatory pathways, expectation appears to bias a shared ACC–PAG circuit toward either inhibitory or facilitatory output.

The human relevance of our findings is enhanced by the reverse translational approach used, building on human pharmacological data. Evidence of human pain as a learning phenomenon continues to build(Seymour 2019), as does evidence for the social modulation of pain(Krahe, Springer et al. 2013, Martin, Tuttle et al. 2014, Mogil 2019, Cho, Deol et al. 2021). The prognosis of pain patients is worsened by sometimes inadvertent nocebo suggestions by clinical providers(Colloca and Finniss 2012), and substantial psychological risk factors such as anxiety, harm avoidance, and pain catastrophizing can be conceptualized as an auto-nocebo process(Corsi and Colloca 2017). However, nocebo effects are generally greater in female subjects (Manai, van Middendorp et al. 2019), whereas we did not observe differences between male and female mice in our nocebo models. This discrepancy may reflect species differences in baseline anxiety levels, hormonal regulation, or variations in experimental paradigms. Addressing these factors through comparative studies involving additional species, and human-like conditioning protocols will help clarify the observed differences and strengthen the translational applicability of our findings.

By isolating CCK-dependent signaling from the ACC to the lPAG as a key mechanism for nocebo hyperalgesia, our findings distinguish negative expectation-driven pain amplification from generalized stress or injury-related hypersensitivity. More broadly, the mouse models described here provide a controlled experimental approach for examining how cognitive and social context shape nociceptive processing, offering a basis for future studies of negative expectation-dependent modulation of pain.

## Supporting information

Poulsonetal_CCKInputs_Supp

## Corresponding authors

Correspondence and requests for materials should be addressed to

## Author contributions

Conceptualization: SJP, JSM, LJM; Methodology: SJP, AS, FTZ, DCB, SAK, LVP, WC, AM, JB, ZS, MF, SG, OBM, MD, AD, LL, RC, LJM; Investigation: SJP, AS, FTZ, DCB, SAK, LVP, WC, AM, JB, ZS, MF, SG, OBM, MD, AD, LL, RC, LJM; Visualization: SJP, JSM, LJM; Funding acquisition: LJM; Supervision: JSM, LJM; Writing: LJM prepared the original draft with SJP, AS, and JSM contributing specific sections.

## Data and materials availability

All data needed to evaluate the conclusions in the paper are present in the paper. Supplementary Information/Source Data are provided with this paper.

## Supplementary Materials

Supplementary Figs. 1 to 19

## Materials and Methods

### Experimental animals

Most experiments used naive, adult CD-1 (ICR:Crl) or C57BL/6 mice aged 6–13 weeks, of both sexes. CD-1 mice were bred in-house at McGill University, originating from breeders obtained from Charles River (St. Constant, QC). C57BL/6 mice were purchased from Charles River (St. Constant, QC) and housed in the University of Toronto Mississauga (UTM) vivarium for at least two weeks prior to starting behavioral testing. Experiments targeting CCK neurons used offspring bred in-house (UTM) from CCK-IRES-Cre mice, which were originally purchased from The Jackson Laboratory (Bar Harbor, ME; Stock #012706). All mice were group-housed in cages of 2–4 on individually ventilated cage racks (Techniplast North America) in a temperature-controlled (22 ± 2°C) room with access to food (Harlan Teklad 8604 for McGill mice; 2019 Teklad for UTM mice) and water *ad libitum*. Both vivaria were maintained under a temperature-controlled environment and operated on a 12:12 h light/dark cycle (lights on at 07:00 h). Mice were allocated randomly to experimental groups and testing was conducted by multiple female experimenters^1^ who were blind to drug treatments. All procedures adhered strictly to Canadian Council on Animal Care regulations and received approval from local animal care and use committees.

### Pharmacological agents

Proglumide (10 mg/kg; Tocris or Sigma Aldrich) was dissolved in saline and administered intraperitoneally (i.p.). LY225910, the CCK-2 antagonist (1 mg/kg, Tocris) was dissolved in 5% DMSO in saline using a sonicator or 5% DMSO and 0.5% Tween-80 in saline and administered intraperitoneally (i.p.). Systemically administered drugs were injected using a volume of 10 ml/kg body weight. In microinfusion experiments, proglumide (250 ng in 250 nl aCSF, Sigma Aldrich) or CCK-sulfated octapeptide (CCK-8S) (50 ng in 250 nl aCSF, Cayman Chemicals) were microinfused through implanted cannulas via calibrated tubing (see below).

### Viral reagents

Viruses were injected at a volume of 200 nl for functional experiments and 100 nl for tracing. AAVretro-CAG-DIO-mNeptune, AAV2/9-hSYN1-SIO-eOPN3-mScarlet, AAV2/9-EF1α – DIO-ChR2(H134R)-mCherry, and AAV2/9-EF1α-DIO-mCherry were purchased from Neurophotonics (Quebec City, QC). AAV2/8-hSYN-DIO-mCherry and AAV-2/8-CamkIIa-GFP were purchased from Addgene (Watertown, MA).

### Pain hypersensitivity conditioning paradigms (Contextual nocebo)

The conditioning paradigms comprised three distinct phases: baseline testing where basal thresholds were measured, the learning session, during which the association between pain (unconditioned stimulus, UCS) and context (conditioned stimulus, CS) was formed, and the testing day, which assessed the response to the presentation of the CS. Two different pain stimuli were used as the UCSs: hind paw incision, or chronic constriction injury, as described below.

Prior to surgery, mechanical thresholds were assessed using the von Frey test (see below). On the learning day, mice underwent either hind paw incision surgery on their left paw to induce post-surgical pain, or chronic constriction injury (CCI) on their left leg, which served as the UCSs. Following surgery, while still under anesthesia, the mice were placed in the conditioning context, consisting of Plexiglas cubicles within a specific laboratory testing room, where they remained for 5 hours, most of which was spent awake. Mice subjected to incision and CCI surgery were returned to the same or different context at 6 or 14 days after surgery, respectively. On test days, mice were retested for pain sensitivity following a 10–90-min habituation period using the manual von Frey test. For baseline and test day measurements, four separate determinations, two per hind paw, were conducted and averaged.

#### Hind paw incision surgery

The hind paw incision surgery was done following the protocol of Cowie and Stucky^2^. Mice were anesthetized by continuous inhalation of isoflurane. A 5-mm incision through the skin and facia was performed on the left hind paw of the mouse. The flexor digitorum brevis muscle was elevated with forceps or blunt dissection probe, and a longitudinal incision was made through the muscle by a scalpel. The wound was closed with two skin sutures.

#### Chronic constriction injury (CCI) surgery

The CCI was performed following the procedure of Bennett and Xie^3^. The sciatic nerve was exposed midway along the thigh using gentle dissection through the biceps femoris muscle. Preceding the trifurcation of the sciatic nerve, approximately 7 mm of nerve tissue was released from surrounding adhesions. Three ligatures (7-0 silicone coated braided silk, Sofsilk) were loosely tied around the nerve, spaced roughly 1 mm apart. The affected nerve segment measured between 4–5 mm in length. Particular attention was paid to tying the ligatures to ensure minimal constriction of the nerve when examined under 40x magnification. The intended level of constriction impeded, but did not fully halt, circulation through the outer blood vessels of the nerve, occasionally eliciting a brief muscle twitch around the exposed area. The incision was subsequently closed in layers. This assay mimics neuropathic pain states in humans, and in our hands causes hypersensitivity lasting more than a month^4^.

### One-in-pain social interaction (Social nocebo)

Same-sex cagemate pairs were placed in an open top/bottom box (inner dimensions: 14.5 cm x 14.5 cm x 30 cm) with three opaque sides and one clear side. Each box was set on top of a clear acrylic platform, which was either placed on a frame for the formalin assay to record video from below or set on top of a lab bench for all other assays. For each experiment, pairs of mice were habituated to the box for 1 h. Mice in the pair were randomly assigned as either observer or demonstrator. Following habituation, the demonstrator was removed briefly, administered an intraplantar hind paw injection of formalin (2.5%, 20 μl; Pain Observer condition), or 0.09% saline as a no-pain control stimulus (No Pain Observer condition). Formalin hind paw injection was selected as the pain stimulus for the Pain Observer condition due to the characteristic pattern of overt nocifensive behaviors that are reliably perceived by observer mice^5,6^. The observer and demonstrator were then allowed to interact for 30 min, a duration selected based on previously published social transfer/emotional contagion models^5–8^. Following the 30 min one-in-pain social interaction, the demonstrator was removed from the box and returned to its home cage. The observer was then either tested for formalin sensitivity (20 μl, 0.5%) intraplantar to the left hind paw or placed in individual Plexiglas cubicles over a metal mesh platform for electronic von Frey testing (see below). For von Frey testing, baseline measurements consisted of five separate determinations for each hind paw, which were averaged. The observer and demonstrator underwent the social nocebo procedure, after which the observer was tested for mechanical sensitivity. In one experiment, mice were tested immediately after, 4 h after, and 24 h after the one-in-pain social interaction. However, most von Frey experiments for the social nocebo paradigm report the average of two measurements per paw starting at 10 min after placing the observer into the Plexiglas cubicle.

#### Quantification of social behavior

In a subset of mice, behaviors during the 30-min one-in-pain social interaction were counted. Behaviors were sampled during the first 10 s of every minute. Data were quantified as the percent positive samples across the 30-min session. In the demonstrator mouse we recorded hind paw licking. In the observer mouse we recorded: (1) attending toward the cagemate including sniffing or grooming, excluding the anogenital region, and (2) propinquity behavior defined as being within 2 cm or less of the demonstrator mouse, but not actively engaging in attending behavior.

### Pain behavior tests

#### Radiant heat paw-withdrawal test

The Hargreaves’ radiant heat paw-withdrawal test was used to assess acute thermal nociception sensitivity in mice^9^. Prior to behavioral testing, mice were individually housed in perforated, transparent Plexiglas enclosures (5 cm by 8.5 cm by 6 cm) situated on a glass floor with a thickness of 3/16^th^ inches and allowed to acclimate for 2 h. A high-intensity beam (IITC model 336; ≈ 45 W) emitted from a projector lamp bulb positioned 6 cm beneath the glass floor was directed at the plantar surface of the mid-hind paw of each mouse at rest. Withdrawal latency of each hind paw was recorded to the nearest 0.1 s.

#### von Frey test

Testing of mechanical sensitivity was performed using either the manual or electronic von Frey test, depending on the laboratory where the experiment was performed. *Manual von Frey:* In contextual nocebo experiments, mice were placed individually in transparent Plexiglas cubicles (5 cm x 8.5 cm x 6 cm) on top of a perforated metal floor and habituated for 2 h before testing. Nylon monofilaments (force range: ≈0.015 to 1.3 g; Touch Test Sensory Evaluator Kit, Stoelting) were firmly applied to the plantar surface of each hind paw of an inactive mouse until they bowed for 0.5 s. The up-down method of Dixon^10^ was used to estimate 50% withdrawal thresholds. *Electronic von Frey:* In social nocebo and cannulation experiments, observers were placed in individual Plexiglas cubicles (10 cm x 5.5 cm x 6.5 cm) over a metal mesh platform and allowed to habituate for 30 min prior to baseline testing. Mice were tested using a Dynamic Plantar Aesthesiometer (Ugo Basile, 37450, Gemonia, Italy) which raised a blunt 0.5-mm metal probe onto the center of the hind paw and gradually applied force (0–10 g ramp over 10 s) until the mouse withdrew its hind paw. The maximal pressure displayed upon withdrawal was recorded.

#### Formalin test

In social nocebo experiments, chemical pain sensitivity in observer mice was assessed using a low concentration formalin challenge (0.5%, 20 μl, intraplantar injection into the left hind paw). This concentration was chosen because it elicits minimal nocifensive behavior while avoiding behavioral ceiling effects that are typically observed at higher concentrations^11^. Following injection, observer mice were returned to the test box, and behavior was video-recorded for 60 min. Videos were scored using a time-sampling procedure in which the presence of licking or biting of the injected paw was recorded during the first 10 s of every min for a total of 60 observations per mouse. We based this procedure on that used by Chesler et al. (2003),^12^ where formalin behavior was quantified using a similar time-sampling procedure that can be seen in Figure 2 of that paper. Based on our experience, this method provides better inter-rater reliability than continuous scoring while remaining sensitive to changes in nocifensive behavior, particularly when using low formalin concentrations. Formalin hind paw injection typically presents a pattern of vigorous licking in the first 5–10 min, deemed the “early phase”, a quiescent period in the next 5–10 min where very little licking occurs, then a “late phase” of sporadic licking that occurs throughout the remainder of the testing period^13,14^. While total time spent licking is often reported in formalin studies, the time-sampling approach used here has been used by our labs across several studies^15–19^. Thus, the formalin data presented throughout the paper represent the percentage of positive samples across the late phase of the test.

#### Hot-plate thermal sensitivity test

The hot-plate test was used to assess whether applying CCK to the dmPAG or lPAG enhanced thermal sensitivity. Mice were tested in a counterbalanced manner on the hot plate 30 min after microinfusion with CCK or aCSF, then 48 h later were retested with the other drug. After microinfusion, mice were immediately moved to the testing room. Mice were then placed singly on a clean hot plate set at 52.5 °C for 45 s. Videos were analyzed using BORIS (Behavioral Observation Research Interactive Software)^20^ for the cumulative number of hind paw flicks and the latency to first hind paw flick, lick, or jump.

### Predator threat–evoked hypersensitivity paradigm

To assess whether CCK signaling contributes to unconditioned stress-evoked pain modulation, mice underwent a single-exposure predator odor conditioning procedure using trimethylthiazoline (TMT). Baseline mechanical sensitivity was first measured using manual von Frey testing. Twenty-four hours later, mice were allowed 5 min of free exploration in a two-chamber apparatus to determine side preference. After an additional 24 h, mice received proglumide (10 mg/kg) or saline 30 min before conditioning. They were first placed in the non-preferred chamber for a 5-min neutral exposure and, 2 h later, exposed for 5 min to filter paper containing 10% TMT. Mechanical sensitivity was reassessed 3 h after TMT using the von Frey test. Contextual memory and defensive behaviour were subsequently evaluated using a conditioned place aversion procedure, followed 24 h later by a 5-min preference test. Manual von Frey thresholds were then measured across the following 21 days. Ten days after the conditioned place aversion test, conditioned freezing was quantified during 5-min re-exposures to both the unpaired and TMT-paired chambers. This delayed assessment was included because we previously demonstrated that blocking corticosterone synthesis prior to initial TMT exposure prevents both unconditioned freezing to TMT and conditioned freezing to the TMT-paired context when mice were returned 10 days later^21^. The TMT paradigm differs from contextual nocebo conditioning in that hypersensitivity is triggered by an unconditioned threat rather than a learned pain-predictive association.

### Stereotaxic surgery and related manipulations

All surgeries were performed under sterile conditions. Mice were anesthetized with isoflurane (4% induction, 2–2.5% maintenance) and body heat maintained on a circulating heated water mat. The scalp was sterilized, shaved, a small incision was made in the skin, the skull leveled using a digital stereotaxic instrument (Harvard Apparatus, Holliston, MA), and burr holes drilled at the respective stereotaxic coordinates. Mice were treated with postsurgical analgesia (20 mg/kg meloxicam) once daily for three days.

#### Cannulations and microinfusion experiments

Mice were cannulated with either a single cannula for drug application to the dmPAG or a double cannula for bilateral drug application to the lPAG. Separate cohorts of mice for both region and drug were used and then tested on either von Frey or hot-plate tests. A single guide cannula with length 3.0 mm and injector cut 0.5 mm below the pedestal (C315GS-5/spc, P1 Technologies, Roanoke, VA) was implanted above the dmPAG (11.7° angle, AP –3.6 mm, ML –0.2 mm, DV –2.55 mm), or a double guide cannula (RWD Life Science, San Diego, CA: guide cannula 62056, center distance 1.4 mm, length 3.0 mm, dummy 62127, injector 62056, both cut 0.5 mm below guide cannula, cap 62525) was implanted above the lPAG (AP –3.85 mm, ML ±0.55 mm, DV –2.65 mm) at 7 weeks of age. Following a 7-day recovery period, mice were tested for baseline von Frey sensitivity and then either underwent contextual nocebo or social nocebo procedures (see above). In all experiments, microinfusions were performed under light isoflurane anesthesia immediately preceding context exposure or the 1-h habituation before social nocebo. In **Fig. 3, d to i**, infusions occurred 30–45 min before pain testing. During infusion, calibrated tubing was used to connect the internal cannula to a microliter syringe (1700 series Hamilton Syringe, 100 μl, Harvard Apparatus, Montreal QC) and an infusion pump (Pump 11 Elite Nanomite, Harvard Apparatus, Montreal QC) was used to infuse a 0.25-μl volume over 5 min (50 nl/min), after which the injector was left in place for an additional 1 min. Mice were allowed to recover in their home cage prior to testing. Cannula placement was verified post hoc by infusing 250 nl of fluorescent muscimol-BODIPY (M23400, Invitrogen) and quantifying the radial spread of fluorescence for both dmPAG (single cannula) and lPAG (bilateral cannula) experiments. The estimated spread radius was approximately 300–350 μm and was comparable across hemispheres in bilateral infusions and similar in magnitude to single-cannula infusions. Mice with placements outside the intended target region (dmPAG or lPAG) were excluded (<5%).

#### Viral infusions

Viruses were infused at a rate of 60 nl/min using a glass capillary micropipette backfilled with mineral oil and attached to a 1700 series gas-tight Hamilton microsyringe. Micropipettes were lowered 0.1 mm below injection site (ACC: AP +1.0 mm, ML ±0.5 mm, DV –0.8 mm; lPAG: AP –3.85 mm, ML ±0.5 mm, DV –2.65 mm) prior to infusion and left in position for 10 min prior to slow removal to prevent backflow. Skin was sutured and the mouse recovered on a heated pad. Viruses were allowed three weeks of expression prior to behavioral testing or tracing.

#### Optogenetic manipulations

CCK-IRES-Cre mice were infused with virus expressing eOPN3, ChR2, or control in the lPAG. In the same surgery session, an optic fiber cannula (200 μm core, 3.0 mm length, R-FOC-BL200C-39NA, RWD Life Science, San Diego, CA) was implanted above the lPAG and secured with dental cement (3M ESPE RelyX Unicem Aplicap Self-Adhesive Universal Resin Cement, 56818). The virus was allowed to express for three weeks. *<u>Contextual nocebo:</u>* For the hind paw incision eOPN3 experiment, viruses were injected 2 weeks prior to baseline, followed by hind paw incision surgery and conditioning to the cubicles. This allowed the virus to express for a total of 3 weeks before mice were tested on test day. In addition, mice were connected to the patch cord (200 nm, Doric, Quebec City, QC) for 10 min at the beginning of the 5 h conditioning period in the contextual nocebo experiments. Then, on the test day mice were tethered and immediately put into the observation box, exposed to that box for 30 min, then put into the von Frey cubicle and allowed to rest for 10 min. They were then tested for mechanical sensitivity using the manual von Frey test. *<u>Social nocebo:</u>* Two days prior to testing in the social nocebo paradigm, pairs were habituated to the box for 1 h. One day prior to testing, pairs were habituated again to the box for 30 min, then the observer mouse was habituated to the patch cord in the box for 10 min. On test day, pairs were habituated for 1 h to the container, then the patch cord was attached to the observer mouse that was allowed to further acclimate for 5 min. *<u>Optical Stimuation for eOPN3:</u>* Light delivery was initiated immediately after placement in the conditioning box for contextual nocebo or following formalin injection in the demonstrator for social nocebo. Optical stimulation consisted of 50 ms pulses at a light power density of 10 mW/mm^2^ of 470 nm light (470 nm, 50 W Collimated LED BLS-LCS-0470-50-22, Oasis Implant, Polyscan3 software, Mightex, Toronto, Canada) delivered once every 2 min to the lPAG of the conditioned mouse or observer for the 30 min duration. Mice were then moved to individual Plexiglas cubicles over a metal mesh platform and tested for mechanical sensitivity. Values reported are the average of two measures on each hind paw. In one experiment, light was delievered in 10-min epochs spaced at 1 h intervals throughout conditioning to minimize tissue heating and phototoxicity during prolonged inhibition. *<u>Optical Stimulation for ChR2:</u>* Mice were tested for baseline mechanical sensitivity. Mice were then habituated to the stimulation box for 30 min during two consecutive days prior to testing with 10 min of habituation to the patch cord each day. On test day, mice were habituated to the box with patch cord for 10 min prior to stimulation. For optogenetic photostimulation of ChR2, light output was 10 mW/mm^2^ delivered at cycles of 5 ms bursts of light on and 45 ms of light off for a frequency stimulation of 20 Hz over a 3 min period. Mice were immediately removed from the stimulation box and placed in von Frey cubicles. In one experiment, mice were tested using manual von Frey and values reported represent the average of two measures on the left hind paw. In another experiment, mice were tested using electronic von Frey with the baseline representing the average of three measures and then subsequent time points (5, 15, 30, 45, 60, 90, and 120 minutes post-stimulation) are a single measure per hind paw.

### Histological procedures, imaging and cell counting

Mice were deeply anesthetized with sodium pentobarbital (100 mg/kg, i.p.) and underwent transcardiac perfusion with 0.1 M phosphate-buffered saline (PBS, pH = 7.4) followed by 4% (w/v) paraformaldehyde in PBS (PFA). Brain tissue was extracted and post-fixed for 24 h in 4% PFA, then cryoprotected for 48 h in 30% (w/v) sucrose. Tissue was embedded in cutting medium (Tissue-Plus O.C.T. Compound, Fisher Scientific Canada) and frozen. Frozen tissue was then sectioned coronally on a cryostat (Cryostar NX50, ThermoFisher Scientific) at 40-μm thickness, then collected in cryoprotectant (30% w/v sucrose, 30% v/v ethylene glycol, 0.1 M PBS) until use.

#### Immunohistochemistry

Sections were thawed, placed in a 12-well plate, then washed three times for 5 min in 0.1 M PBS (pH = 7.4). Sections were blocked (5% normal goat serum, 0.3% Triton X-100, 0.1 M PBS) for 2 h at room temperature (RT). The block was removed, and sections were then incubated in primary antibody at 4°C. For c-Fos studies, a rabbit monoclonal antibody (rabbit α c-Fos 1:10,000 for 48 h, Synaptic Systems 226008, Göttingen, Germany) was used. In the retrograde tracing experiment, a monoclonal NeuN antibody (1:2,000 overnight, EMD Millipore MABN140, Darmstadt, Germany) or a polyclonal EMX1 antibody (1:50 overnight, Invitrogen, PA5-35373, Carlsbad, CA), were used as general or excitatory neuronal markers, respectively. Following primary antibody incubation, sections were washed three times for 5 min in PBS, then incubated with secondary antibody (*c-Fos experiments*: goat anti-rabbit, 1:500, 2 h at RT, Vector Laboratories DI-1649-1.5, Newark, CA; *NeuN and EMX1 experiments:* goat anti-rabbit Alexa Fluor 488, 1:500, 2 h at RT, Invitrogen A11008, Waltham, MA). Sections were then washed three times for 5 min in PBS, mounted on Superfrost slides and dipped in DAPI solution (1:50,000 in 0.1 M PBS, 5 min, D9542, Sigma Aldrich Canada, Oakville, ON) and coverslipped with mounting medium (Fluoromount-G, SouthernBiotech, Birmingham, AL).

#### Imaging and analysis of c-Fos

Digital images of coronal brain sections were taken at 4X (1.6125 μm/pixel, Biotek Cytation 5, Agilent, Santa Clara, CA) for the social nocebo paradigm or 10X (Olympus VS200 slide scanner, Evident Corporation, Tokyo, Japan) for the contextual nocebo paradigm using filters for both DAPI and Cy5. QuPath v0.3.2^22^ was used along with the Allen Mouse Brain Reference Atlas Version 2 (Allen Institute for Brain Science, 2011) and schematic of the columns of the rat PAG^23^ to draw regions of interest (ROI) around the anterior, intermediate, and posterior PAG. In addition, we used the DAPI channel to visualize changes in cell density that distinguish columnar organization of the PAG. c-Fos positive cells were determined as high grayscale signal, oval-shaped nuclei and counted using the QuPath cell detection feature followed by manual addition and subtraction of cells by researchers blinded to condition. c-Fos-positive regions were quantified and reported as number of cells/mm^2^, and at least three sections between the bregma range were averaged for each ROI. An unbiased approach was taken such that sections from the anterior, intermediate, and posterior PAG as follows: Anterior PAG: between bregma –2.3 and –2.6. Intermediate PAG: between bregma –3.6 and – 3.9. Posterior PAG: between bregma –4.7 and –4.9. For cortical analyses, c-Fos quantification was performed separately across cortical layers (L1, L2/3, L5, L6) and for retrogradely labeled neurons to distinguish general cortical activation from activation within projection-defined populations.

#### Retrograde tracer and NeuN/EMX1 analysis

The density of CCK-positive (tagged by virus) to NeuN/EMX1-positive cells was compared between selected brain regions (average of 3 slices per mouse from 2 mice for EMX1, average of 3 slices per mouse from 3 mice for NeuN). Digital images of coronal brain sections were taken at taken at 10X (Olympus VS200 slide scanner) using filters for DAPI, GFP (NeuN/EMX1) and mCherry (retrograde virus). QuPath was used along with the Allen Mouse Brain Reference Atlas Version 2 (Allen Institute for Brain Science, 2011) to draw regions of interest (ROI) in the ACC (AP +0.9 to +1.15, ACAd and ACAv), insula (AP +0.9 to +1.15, AId and AIv), motor cortex (AP –1.25 to –0.95, MOs and MOp), retrosplenial cortex (RSPd and RSPv combined), parietal cortex (AP –2.35 to –2.15, PTLp), and lPAG (AP –3.9 to –3.7). Quantification boundaries were drawn around layer V in each cortical region because retrogradely labeled cells were most abundant in this layer. NeuN-positive cells were determined as high grayscale signal, oval-shaped nuclei and counted using the QuPath cell detection feature followed by manual addition and subtraction of cells, whereas mCherry-positive (CCK^+^) cells were quantified using the manual point counting feature. CCK cell density was calculated as a percentage of the mCherry-positive cells within in the ROI (mm^2^) relative to the total number of NeuN-positive cells within the ROI. Density counts were averaged across three consecutive coronal slices for each ROI.

#### RNAscope

Tissue was collected and cut at 40 μm free-floating and then mounted on Superfrost Plus Gold slides. The RNAscope Multiplex Fluorescent V2 Assay (Advanced Cell Diagnostics (ACD), Inc., Newark, CA) was performed according to manufacturer instructions (UM 323100) using the fixed-frozen tissue sample pretreatment and RNAscope Protease III. Tissue was probed for *Cckbr* (CCK-2 receptor, ACD probe cat no. 439121) and *Fos* (ACD probe cat no. 316921) mRNA and respective 570 nm and 690 nm opal dyes (Akoya Biosciences). Slides were coverslipped with ProLong Gold mounting media (Invitrogen, Waltham, MA) and imaged within one week at 20X (Olympus VS200 slide scanner, Evident Corporation, Tokyo, Japan). We report active cells that express *Cckbr* in the social nocebo and contextual nocebo paradigms as the percent of *Fos*-positive cells (distinct 690 nm-positive ovals) that also express *Cckbr* (at least three 570 nm dots within a *Fos*-positive cell boundary). Transcriptomic reference data (see Supplementary Fig. 16) indicated that *Cckbr* expression in the PAG is predominantly neuronal, with most expressing glutamatergic markers, and therefore additional neuronal marker co-staining was not performed.

#### Analysis of publicly available transcriptomic datasets

To provide molecular context for Cckbr-expressing neurons in the PAG, we analyzed publicly available whole-brain single-cell and spatial transcriptomic datasets from the Allen Brain Cell Atlas^24^. Neuronal subclasses within the PAG were identified using the reference taxonomy provided by the Allen Institute, and expression of *Cckbr* was quantified across neuronal populations and along the rostro-caudal axis. Co-expression analyses were performed for genes associated with neuromodulatory and pain-related signaling pathways. These analyses were used to determine the distribution of Cckbr across molecularly defined PAG neuronal populations rather than to define cell identity within individual experimental animals.

#### Verification of viral injections

For verification of viral injections, sections were processed as described above. Injection location was considered accurate when expression of the virus was limited to the boundaries of the target brain region. When visualization of the virus was not possible or spread of the virus was beyond the designated target, data were not included. This occurred in less than 5% of the injected mice.

### Statistical analyses

Statistical analyses were performed in IBM SPSS (v28, IBM Corp.) using an α level of 0.05. Depending on the design, data were analyzed using one-way ANOVA, repeated-measures ANOVA, or paired t-tests. Statistical details for main figures are provided in Supplementary File S1 and for supplementary figures in the corresponding legends. When ANOVAs were significant, Šidák or Tukey multiple-comparison tests were applied as appropriate. For box plots, the box represents the interquartile range (Q1–Q3), the center line denotes the median, and whiskers indicate the minimum and maximum values. Graphical figures were generated using GraphPad Prism (v9), Adobe Illustrator (v27.7), and templates adapted from BioRender (2023).

