## Supplementary material for "Cholecystokinin input from the anterior cingulate cortex to the lateral periaqueductal gray mediates nocebo pain behavior in mice": Poulsonetal_CCKInputs_Supp

### Supplementary Figures and Legends

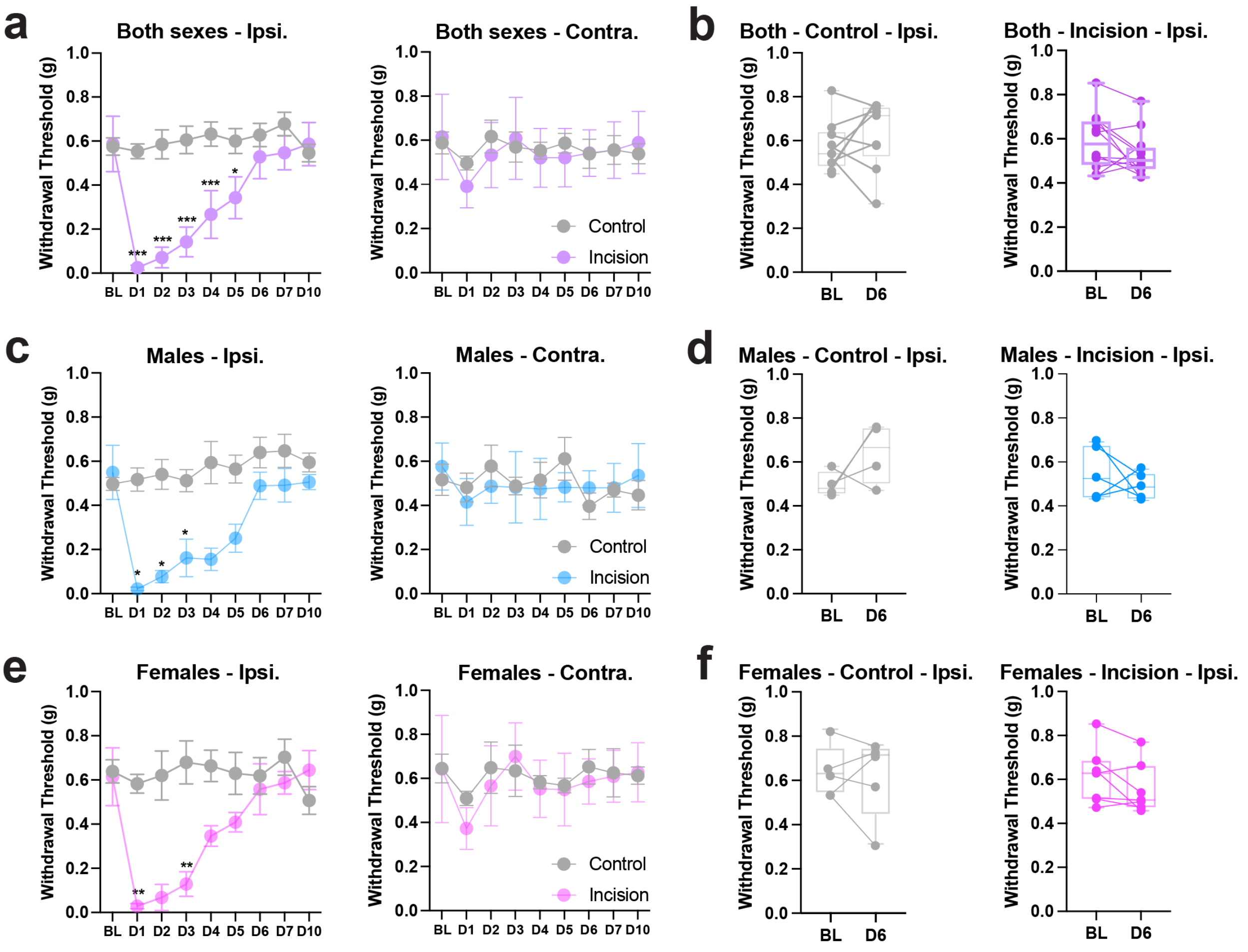


**Supplementary Fig. 1.** Plantar incision produces transient ipsilateral mechanical hypersensitivity that resolves by day 6 in both sexes. **a** Ipsilateral withdrawal thresholds (combined sexes) showed a significant time × surgery interaction (two-way ANOVA, $F(4.98,94.64) = 18.64$, $P < 0.001$). Post hoc Šidák comparisons revealed reduced thresholds on days 1–5 after incision with no difference from baseline at day 6. Contralateral thresholds showed no time × surgery interaction (two-way ANOVA, $F(5.73,108.8) = 0.81$, $p = 0.56$). **b** Paired t-tests (baseline vs. day 6) confirm this recovery with both control and incision groups showing similar thresholds relative to baseline at day 6, indicating full resolution of incision-evoked mechanical hypersensitivity (*control*, $t(8) = 0.85$, $p = 0.41$); *incision* ($t(11)= 1.78$, $p = 0.10$). **c** Males show the same pattern with a clear ipsilateral decrease from days 1–3 followed by recovery (*ipsilateral,* $F(3.358,24.77)=8.37$, $p < 0.001$); *contralateral*, $F(3.81,26.68)=0.89$, $p = 0.48$). **d** Paired t-tests (baseline vs. day 6) confirm this recovery with both control and incision groups showing similar thresholds relative to baseline at day 6, indicating full resolution of incision-evoked mechanical hypersensitivity (*control*, $t(3) = 1.45$, $p = 0.24$); *incision* ($t(4)= 0.87$, $p = 0.43$). **e** Females show the same pattern with a clear ipsilateral decrease on days 1 and 3 followed by recovery (*ipsilateral,* $F(3.71,37.13)=12.12$, $p < 0.001$); *contralateral*, $F(4.976,49.76)=0.89$, $p = 0.41$). **f** Paired t-tests (baseline vs. day 6) confirm this recovery with both control and incision groups showing similar thresholds relative to baseline at day 6, indicating full resolution of incision-evoked mechanical hypersensitivity (*control*, $t(4) = 0.29$, $p = 0.78$); *incision* ($t(6)= 1.76$, $p = 0.12$). Data are mean ± SEM. *$p < 0.05$; **$p < 0.01$; ***$p < 0.01$ from Šidák-corrected post hoc comparisons.

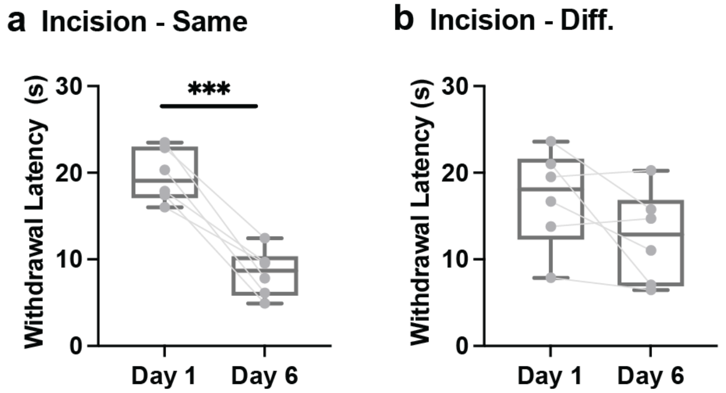


**Supplementary Fig. 2. Thermal withdrawal latencies following contextual nocebo conditioning. a** Mice returned to the same context (n = 6 mice) showed reduced withdrawal latency relative to Day 1. **b** Mice tested in a different context (n = 6 mice) do not show a change in withdrawal latency relative to Day 1. Analysis of thermal sensitivity 6 days after hind paw incision conditioning revealed a significant context × time interaction (two-way ANOVA, $F(1,10) = 5.83$, $p = 0.036$). Box plots show the median and interquartile range, with dots and connecting lines representing paired measurements from individual mice. ***$p < 0.001$ from Šidák-corrected post hoc comparisons.

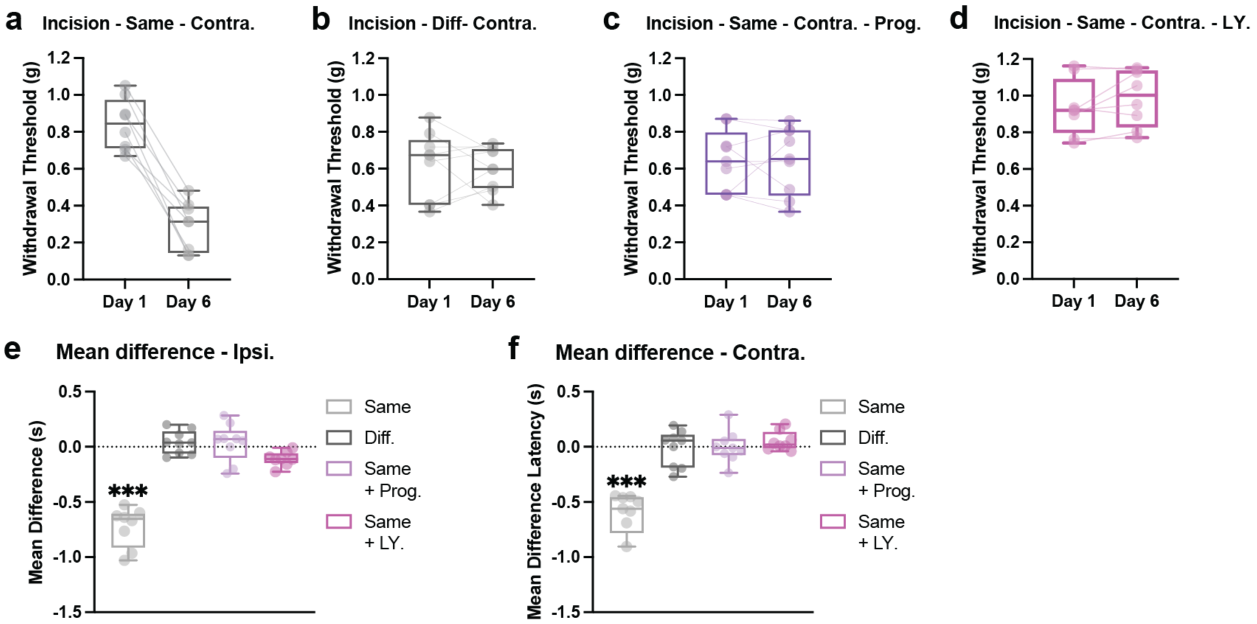


**Supplementary Fig. 3. Contralateral hind paw sensitivity during contextual nocebo conditioning following unilateral incision.** Mechanical withdrawal thresholds (manual von Frey) were measured in the paw opposite the incision before (Day 1) and after conditioning (Day 6). Mice tested in the same context (**a,** n = 8 mice) showed reduced thresholds, whereas mice tested in a different context (**b,** n = 9 mice) did not. Systemic administration of proglumide (**c**, n = 9 mice) or the selective CCK-2 receptor antagonist LY 225910 (**d**, n = 8 mice) prevented the decrease in withdrawal thresholds (two-way ANOVA; condition × time interaction, $F(3,30)=32.31$, $p<0.0001$). **e** Mean change in withdrawal threshold (Day 6 − Day 1) is shown for ipsilateral measurements corresponding to Fig. 1b–e (one-way ANOVA, $F(3,30)=56.92$, $p<0.001$). **f** Mean change in withdrawal threshold for contralateral measurements from panels a–d (one-way ANOVA, $F(3,30)=34.98$, $p<0.0001$). Box plots show the median and interquartile range, with dots and connecting lines representing paired measurements from individual mice. ***$p<0.001$ from Šidák-corrected post hoc comparisons.

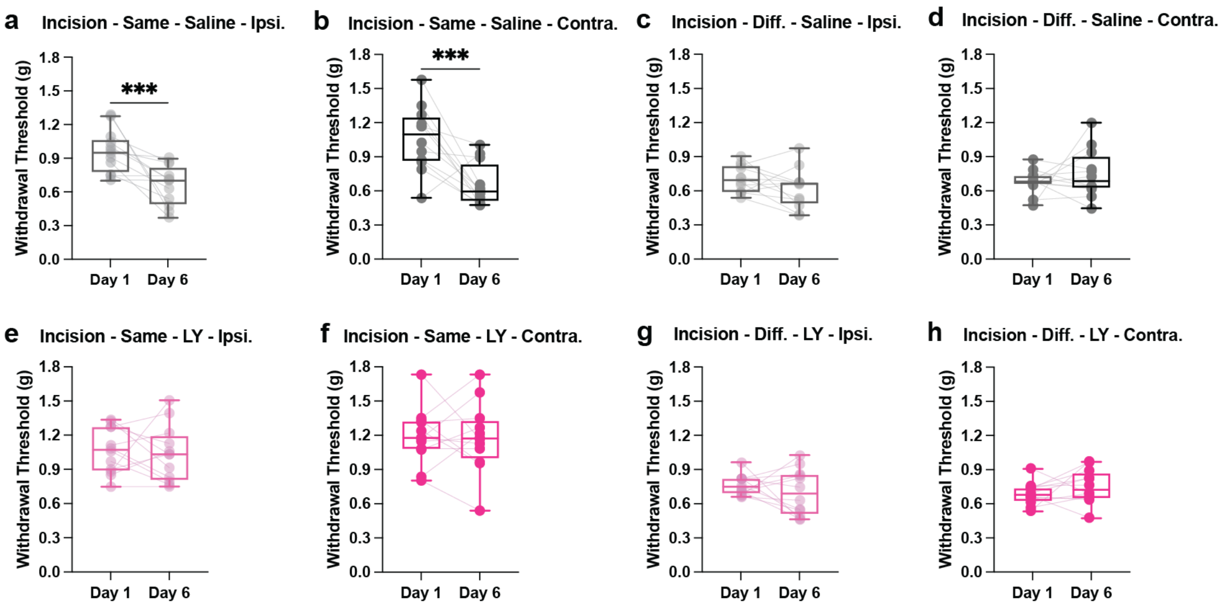


**Supplementary Fig. 4. Bilateral sensitivity following unilateral hind paw incision conditioning is blocked by LY 255910.** Mechanical withdrawal thresholds sensitivity is present in the ipsilateral (**a**) and contralateral (**b**) hind paws of mice returned to the same context (n = 20 mice) following unilateral hind paw incision conditioning. Mice do not show changes in mechanical sensitivity following hind paw incision conditioning when placed in a different context for the ipsilateral (**c**) or contralateral (**d**) hind paws (n = 20 mice). Treatment with the CCK-2 receptor antagonist, LY 225910 blocks mechanical pain sensitivity following hind paw incision conditioning in the ipsilateral (**e**) and contralateral (**f**) hind paws (n = 20 mice). LY 255910 does not change mechanical sensitivity for ipsilateral (**g**) or contralateral (**h**) hind paws when mice are placed in a different context (n = 20 mice). Statistical tests include four-way mixed analysis of variance (ANOVA; context × drug × paw × day) with the three-way interaction of context × drug × day representing the highest order statistically significant effect ($F(1,44)=6.74$, $p=0.013$). Box plots show the median and interquartile range, with dots and connecting lines representing paired measurements from individual mice. ***$p < 0.001$ for post hoc comparisons. ***$p<0.001$ from Šidák-corrected post hoc comparisons.

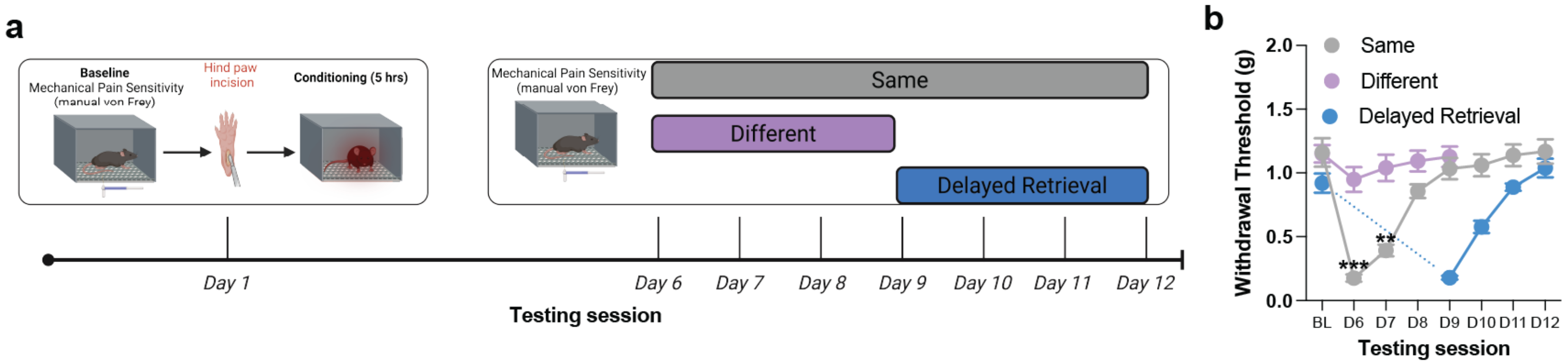


**Supplementary Figure 5. Extinction of context-dependent pain hypersensitivity after hind paw incision. a** Timeline and experimental design showing extinction experiment with repeated testing. Mice received a unilateral hind paw incision on Day 1 paired with a specific context. After incision-evoked hypersensitivity had resolved, mice were returned to the same context from Days 6–12 to assess extinction of conditioned hypersensitivity. Mice tested in a different context were assessed only through Day 9, as hypersensitivity did not develop. In a separate cohort, initial testing was delayed until Day 9 to determine whether hypersensitivity could still be expressed at this later time point (delayed retrieval). **b** Mice returned to the same context (n = 8 mice) produced hypersensitivity on Days 6–7 relative to the different context (n = 8 mice), which progressively resolved with repeated testing, whereas thresholds in the different context remained stable (two-way ANOVA, time × context interaction: $F(2.54, 35.61) = 17.74$, $p < 0.0001$). Mice in the delayed retrieval group (n = 8 mice) revealed hypersensitivity at Day 9–10 relative to mice tested in the same context starting on day 6 (two-way ANOVA, time × context interaction: $F(3.27,45.84) = 17.09$, $p < 0.0001$). Data are shown as mean ± SEM. $**p < 0.01$; $***p < 0.001$ from Šidák-corrected post hoc comparisons.

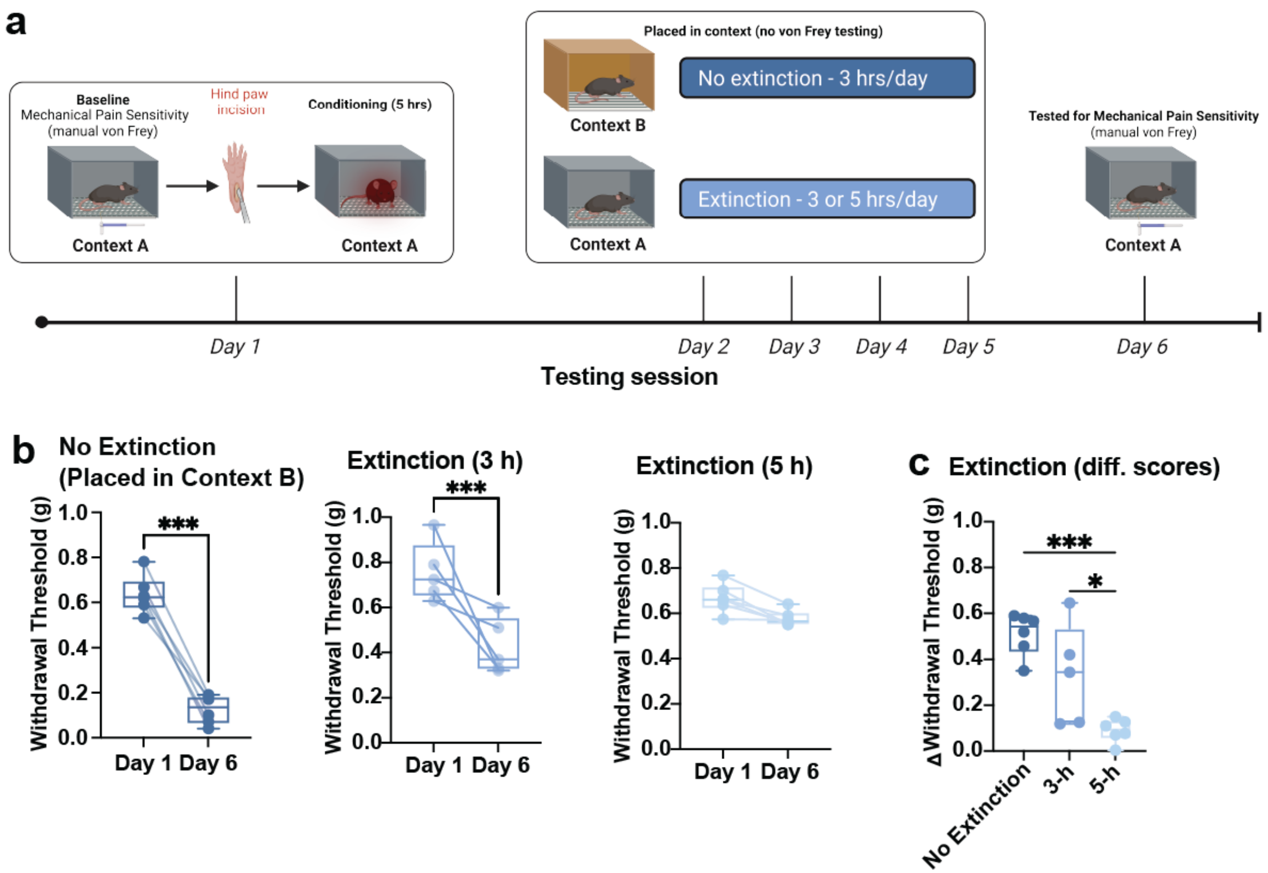


**Supplementary Fig. 6. Repeated passive context exposure reduces conditioned hypersensitivity. a** Timeline and experimental design showing repeated context exposure without testing. Mice were first assessed for mechanical sensitivity in Context A, then received a unilateral hind paw incision and were returned to Context A on Day 1 for a 5-hour conditioning session. On Days 2–5, mice were placed either back into Context A for 3 or 5 h per day or into Context B. Mechanical testing was not performed during these sessions to determine whether context re-exposure alone was sufficient to extinguish conditioned hypersensitivity. On Day 6, mice were returned to Context A for von Frey testing. **b** Mice placed in a different context (n = 6 mice) remained hypersensitive on Day 6, whereas re-exposure to the original context reduced hypersensitivity in a duration-dependent manner (*3 h group*, n = 5 mice, *5 h group*, n = 6 mice; two-way ANOVA, context × time interaction, $F(2,14)=14.92=14.92$, $p<0.001$). Post hoc comparisons showed hypersensitivity in the different-context group, a partial reduction after 3-hour exposure, and no remaining difference after 5-hour exposure ($p<0.001$). **c** Change in withdrawal thresholds (Day 6 – Day 1) differed across groups. Prolonged re-exposure to the conditioning context (5 h/day)

abolished hypersensitivity, shorter re-exposure (3 h/day) produced a partial reduction, whereas exposure to a different context had no effect (one-way ANOVA, $F(2,14)=14.92$, $p<0.0001$). Box plots show the median and interquartile range, with dots and connecting lines representing paired measurements from individual mice. *$p<0.05$, ***$p<0.001$ from Šidák-corrected post hoc comparisons.

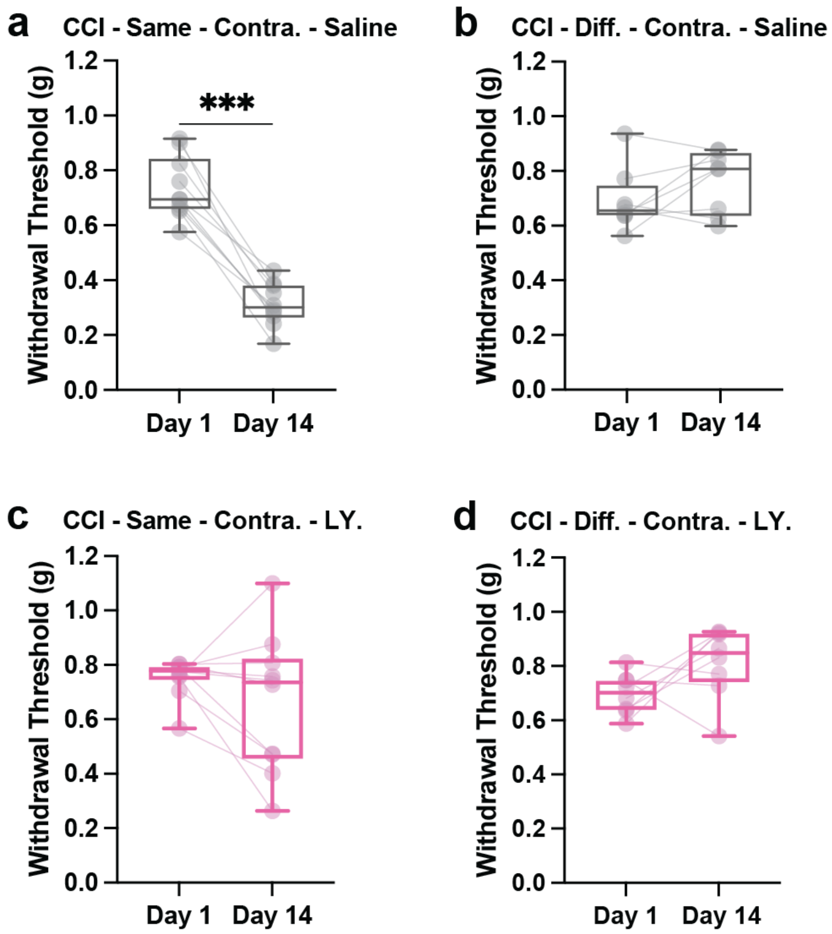


**Supplementary Fig 7. Contralateral hind paw sensitivity during contextual nocebo conditioning following unilateral CCI.** Mechanical withdrawal thresholds (manual von Frey) were measured in the paw opposite the nerve injury before (Day 1) and after conditioning (Day 14). **a** Mice tested in the same context (n = 10) showed reduced thresholds for the contralateral (right paw). **b** Mice tested in a different context (n = 8) do not show changes in withdrawal threshold for the contralateral paw. **c** Systemic administration of the selective CCK-2 receptor antagonist LY225910 prevents the decrease in withdrawal thresholds for the contralateral paw for mice returned to the same context (n = 10 mice). **d** LY 225910 contralateral paw data for mice tested in a different context (n = 8). Statistical tests include three-way ANOVA (context × drug × day interaction; $F(1,32) = 6.00$, $p<0.05$). Box plots show the median and interquartile range, with dots and connecting lines representing paired measurements from individual mice. ***$p < 0.001$ from Bonferroni-corrected paired t-tests.

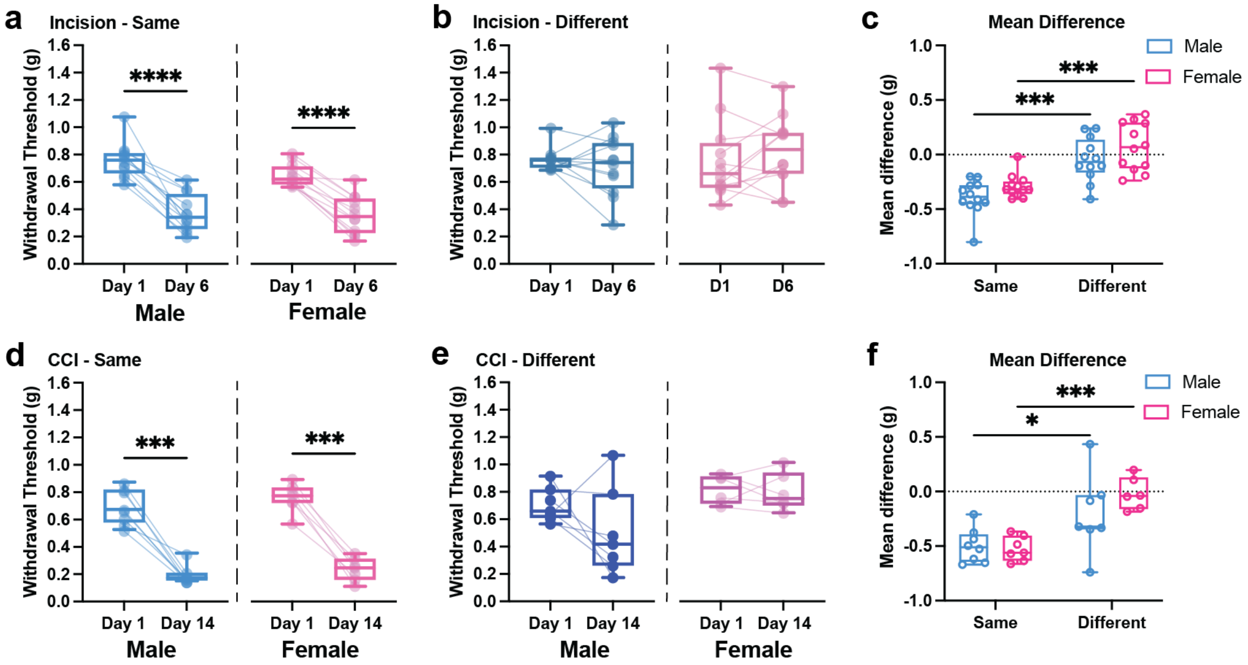


**Supplementary Fig 8. No sex difference in hind paw incision- or CCI-induced conditioned sensitivity.** Independent cohorts of male and female mice were tested to assess sex differences in the contextual nocebo models. **a** Male and female mice that underwent hind paw conditioning show mechanical sensitivity when returned to the same context 6 days after the conditioning session (n = 12 mice/sex). **b** Male and female mice that underwent hind paw conditioning do not show mechanical sensitivity when placed in a different context 6 days after the conditioning session (n = 12 mice/sex). **c** Mean difference scores (Day 6 – Day 1) for male and female mice following hind paw incision conditioning and returned to the same or different context. **d** Male (n = 8) and female (n = 7) mice that underwent chronic constriction injury (CCI) conditioning show mechanical sensitivity when returned to the same context 14 days after the conditioning session. **e** Male (n = 7) and female (n = 6) mice that underwent CCI conditioning do not show mechanical sensitivity when returned to the same context 14 days after the conditioning session. **f** Mean difference scores (Day 14 – Day 1) for male and female mice following CCI conditioning and returned to the same or different context. For both injury-induced conditioning models, the three-way ANOVA (context × sex × day) indicates a significant two-way interaction between context and day (hind paw incision: $F(1,12)=12.913$, $p=0.004$; CCI: $F(1,24)=23.357$, $p<0.001$), but no significant three-way interaction indicating that male and female mice respond similarly to these conditioning sessions (sex × context × day); hind paw: $F(1,12)=0.000$, $p=0.996$; CCI: $F(1,24)=1.641$, $p=0.212$). Two-way ANOVA (sex × room) indicates a significant main effect of room for both conditioning models for main difference scores (hind paw incision: $F(1,12)=12.89$, $p=0.003$; CCI: $F(1,24)=23.53$, $p<0.0001$). Box plots show the median and interquartile range, with dots and connecting lines representing paired measurements from individual mice. *p<0.05, **<p<0.01, ***p<0.001 from Šidák-corrected post hoc comparisons.

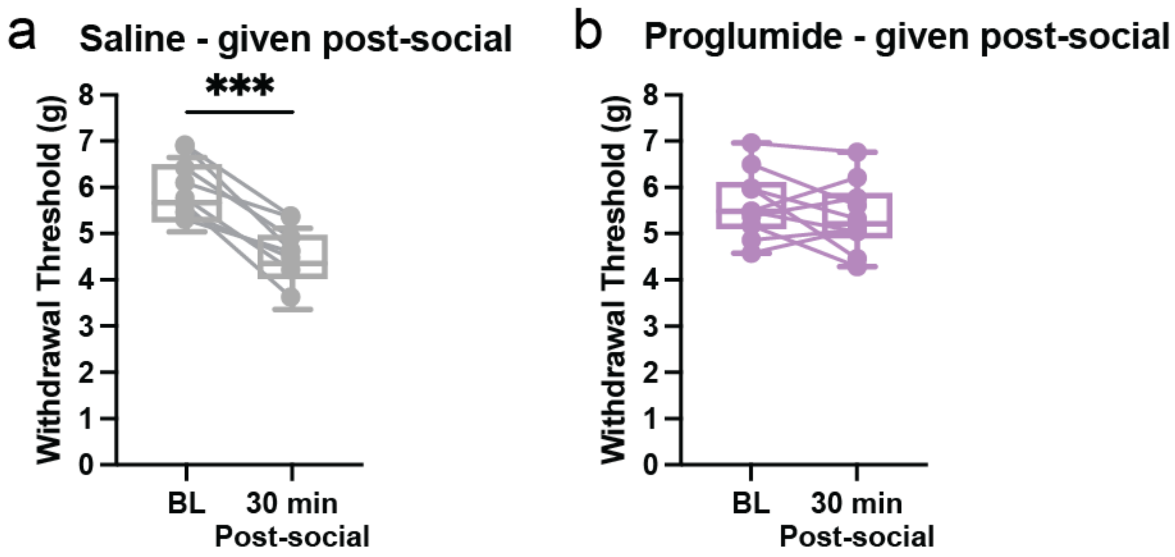


**Supplementary Figure 9. Proglumide administered after social interaction prevents social nocebo–induced mechanical hypersensitivity.** Withdrawal thresholds measured using the electronic von Frey test at baseline (BL) and 30 min after the social interaction. **a** Mice receiving saline (n = 8) after the social interaction showed a significant reduction in withdrawal thresholds at 30 min, indicating increased mechanical sensitivity. **b** Mice treated with proglumide (n = 10) after the social interaction do not show a reduction in withdrawal thresholds. Statistical tests include two-way (drug × time interaction, $F(1,16)=13.12$, $p<0.0001$). Box plots show the median and interquartile range, with dots and connecting lines representing paired measurements from individual mice. ***$p<0.001$ from Šidák-corrected post hoc comparisons.

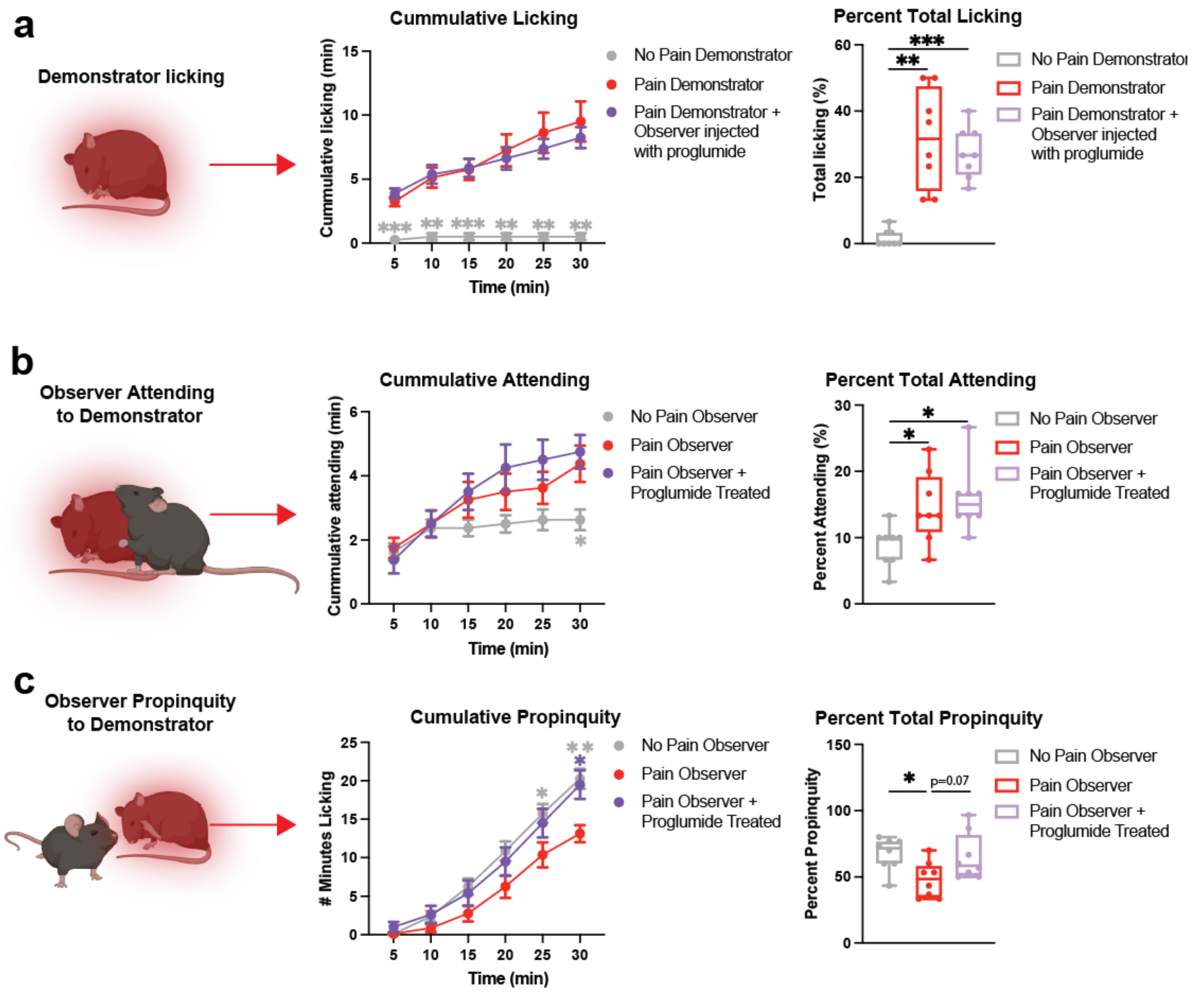


**Supplementary Fig 10. Pain and social behavior related to social nocebo model. a** Paw licking in Pain Demonstrator mice following formalin injection. Pain Demonstrators (n = 8) showed greater licking during the interaction than No Pain Demonstrators (n = 8), and proglumide (n = 8) did not alter licking behaviour. **b** Attending behaviour (sniffing or grooming, excluding anogenital contact). Pain Observers directed more attending behaviour toward Pain Demonstrators than toward No Pain Demonstrators, with no effect of proglumide. **c** Social propinquity (passive proximity $<1$ cm without active interaction). Time course data were analyzed using a two-way ANOVA (a, *Licking*; time × condition interaction, $F(3.09,32.42) = 6.82$, $p = 0.001$; b, *Attending*; time × condition interaction, $F(4.33,45.5) = 4.426$, $p = 0.003$; c, Propinquity; time × condition interaction, $F(4.33,45.5) = 4.43$, $p = 0.003$), while box plots were analyzed using one-way ANOVA (a, *Licking*; $F(2,21) = 22.25$, $p<0.0001$; b, *Attending*; $F(2,21) = 5.51$, $p = 0.012$; c, *Propinquity*; $F(2,21)=4.26$, $p=0.04$). *$p < 0.05$; **$p < 0.01$; ***$p < 0.001$ from Tukey's multiple-comparisons tests. Line graph shows cumulative behavior over a 30-minute social interaction shown as mean ± SEM. Box plots show the median and interquartile range, with dots and connecting lines representing paired measurements from individual mice. Asterisks indicate comparisons to the Pain Observer group, with colour matching the group being compared. *$p<0.05$, **$p<0.01$, ***$p<0.001$ from Šidák-corrected post hoc comparisons.

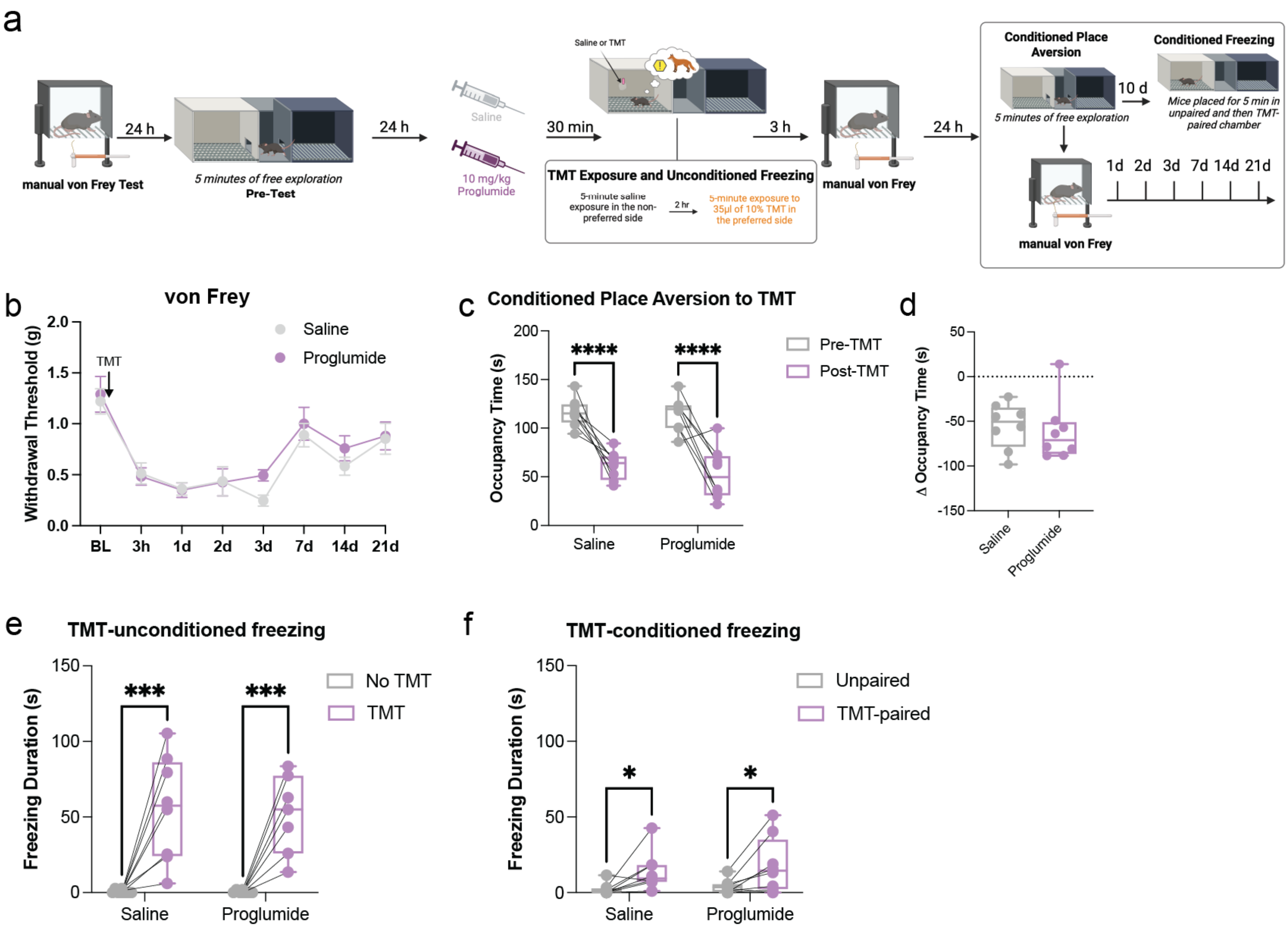

a
24 h
manual von Frey Test
5 minutes of free exploration
Pre-Test
24 h
Saline
10 mg/kg
Proglumide
30 min
Saline or TMT
TMT Exposure and Unconditioned Freezing
5-minute saline exposure in the non-preferred side
2 hr
5-minute exposure to 35µl of 10% TMT in the preferred side
3 h
manual von Frey
24 h
Conditioned Place Aversion
5 minutes of free exploration
10 d
Conditioned Freezing
Mice placed for 5 min in unpaired and then TMT-paired chamber
manual von Frey
1d 2d 3d 7d 14d 21d
b
von Frey
Withdrawal Threshold (g)
TMT
Saline
Proglumide
BL 3h 1d 2d 3d 7d 14d 21d
c
Conditioned Place Aversion to TMT
Occupancy Time (s)
****
****
Pre-TMT
Post-TMT
Saline
Proglumide
d
Δ Occupancy Time (s)
Saline
Proglumide
e
TMT-unconditioned freezing
Freezing Duration (s)
***
***
No TMT
TMT
Saline
Proglumide
f
TMT-conditioned freezing
Freezing Duration (s)
*
*
Unpaired
TMT-paired
Saline
Proglumide

**Supplementary Fig. 11. CCK receptor blockade does not alter predator-threat–evoked nociception or defensive behaviour. a** Experimental timeline for predator-odor (TMT) exposure followed by conditioned place aversion testing, von Frey and re-exposure to the conditioning chambers. Mice received saline or proglumide prior to behavioral assessment. **b** Mechanical withdrawal thresholds across days following TMT exposure showed TMT-evoked hypersensitivity that was not altered by proglumide (n = 8 mice/group; two-way ANOVA, time × drug, p = 0.40.). **c** Mice spent less time in the TMT-paired chamber during the post-test compared to the pre-test regardless of saline or proglumide treatment (n = 8 mice/group; two-way ANOVA, main effect chamber, $F(1,14)=60.64$, $p<0.0001$; no main effect of drug and no context × drug interaction, all p other >0.42). **d** Change in occupancy time between saline and proglumide treatment, calculated within subjects as the difference in time spent in the TMT-paired context ($t(14)=0.42$, $p=0.68$). **e** TMT exposure significantly increased unconditioned freezing compared to no-TMT trials, with no effect of proglumide treatment (two-way ANOVA, main effect time, $F(1,14)=41.02$, $p<0.0001$; no main effect of drug and no context × drug interaction, all other p >0.52). **f** Conditioned freezing to TMT was significantly greater when mice were re-exposed to the TMT-paired context 10 days after initial exposure compared to the unpaired context, with no effect of proglumide treatment (two-way repeated-measures ANOVA, main effect of context, $F(1,14) = 13.37$, $p = 0.003$; no main effect of drug and no context × drug interaction, all other $p > 0.52$). von Frey data is shown as mean ± SEM. Box plots show the median and interquartile range, with dots representing individual mice. *$p < 0.05$; ***$p < 0.001$; p<0.0001 from Šidák-corrected post hoc comparisons.

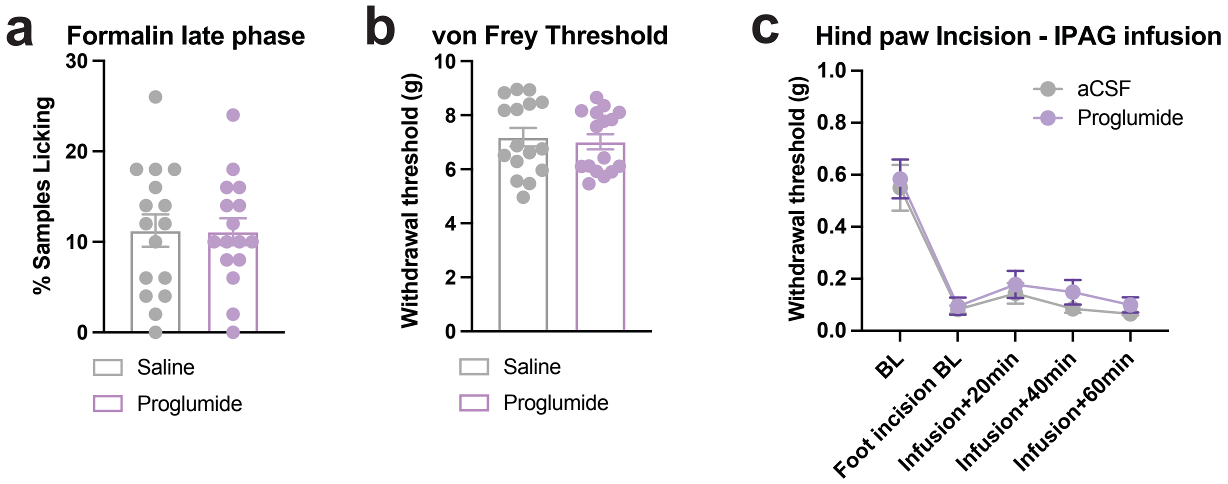


**Supplementary Fig. 12. Proglumide does not alter baseline nociceptive responses or acute injury–evoked pain. a** Late-phase formalin-evoked licking behavior was not altered by systemic proglumide compared with saline controls (*independent t-test;* $t(30)=0.05$, $p=0.96$). **b** Baseline mechanical withdrawal thresholds measured using the electronic von Frey test were comparable between saline- and proglumide-treated mice (*von Frey*, $t(15)=1.03$, $p=0.32$). **c** Following hind paw incision, intra-lPAG infusion of proglumide did not alter the time course or magnitude of incision-evoked mechanical hypersensitivity compared with aCSF controls (two-way ANOVA; *time x drug interaction*, $p = 0.41$). Data are shown as mean ± SEM.

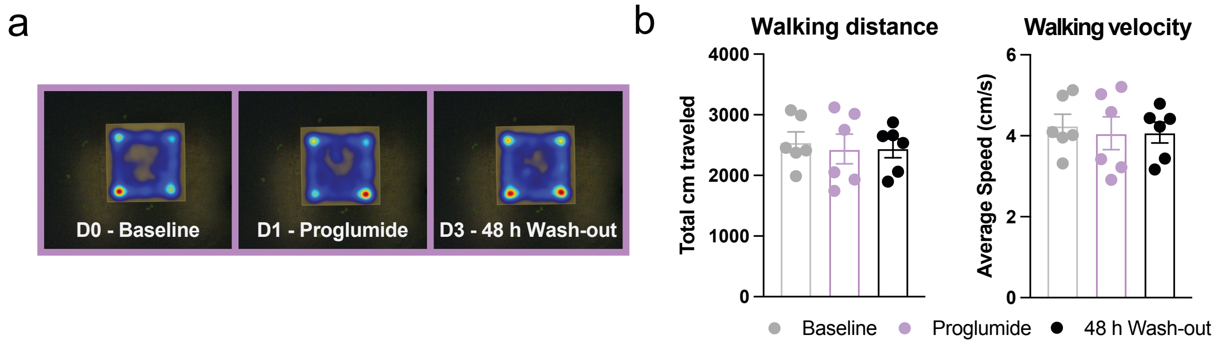


**Supplementary Fig. 13. Proglumide application to the lateral PAG does not influence motor responses in the open field. a** Representative heat maps of mouse activity during baseline (Day 0, D0), 90 min following proglumide microinfusion (Day 1, D1) to the lateral PAG and 48 hr following proglumide infusion (Day 3, D3). **b** Bar graphs show total distance traveled (left) and average speed (right) of experimental groups. One-way ANOVA revealed no statistically significant influence of proglumide on distance traveled (left, $F(2,15) = 0.104$, $p = 0.902$) or velocity (right, $F(2,15) = 0.105$, $p = 0.901$) compared to baseline when applied to the lPAG in the open field test (n = 6 mice). Data are shown as mean ± SEM.

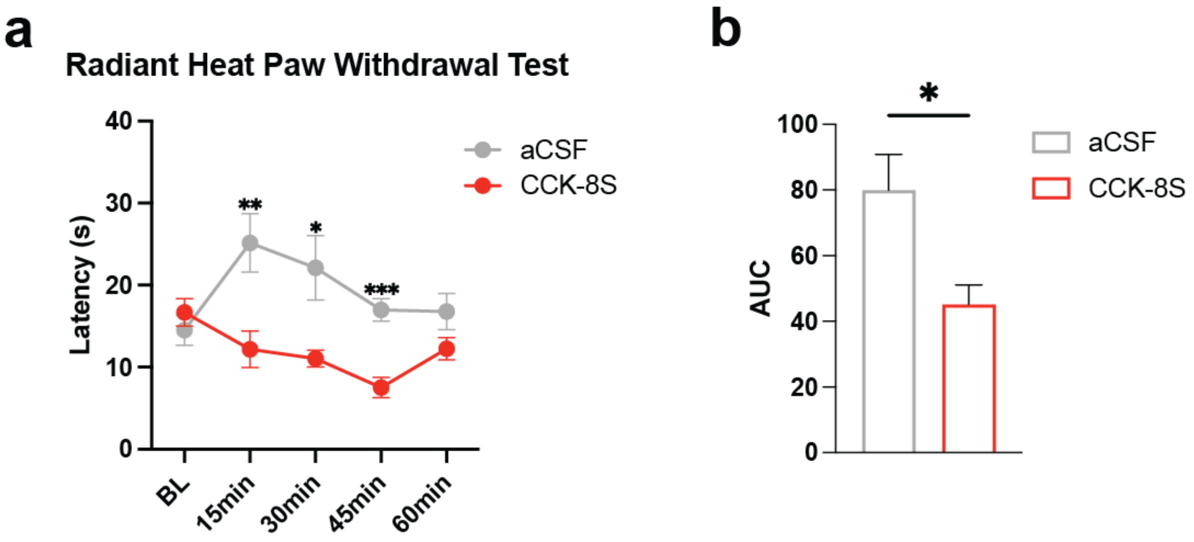


**Supplementary Fig. 14. Application of CCK to the lPAG increases thermal pain responses on the Hargreaves test. a** Withdrawal latency across time following microinfusion of CCK or aCSF into the lPAG (n = 7 mice/group). CCK-treated mice showed reduced withdrawal latency relative to aCSF controls at 15, 30, and 45 min, with no difference at baseline or 60 min. **b** Area under the curve (AUC) analysis corresponding to panel a demonstrating overall reduced latency in CCK-treated mice relative to aCSF controls. Time course data were analyzed using two-way ANOVA (time × drug; $F(4,48)=3.37$, $p=0.016$) followed by Bonferroni-corrected pairwise comparisons (baseline $p=0.396$; 15 min $p=0.01$; 30 min $p=0.02$; 45 min $p<0.001$; 60 min $p=0.110$). AUC values were compared using an independent t-test ($t(12)=2.78$, $p=0.02$). Data are shown as mean ± SEM. *$p<0.05$; **$p<0.01$; ***$p<0.001$.

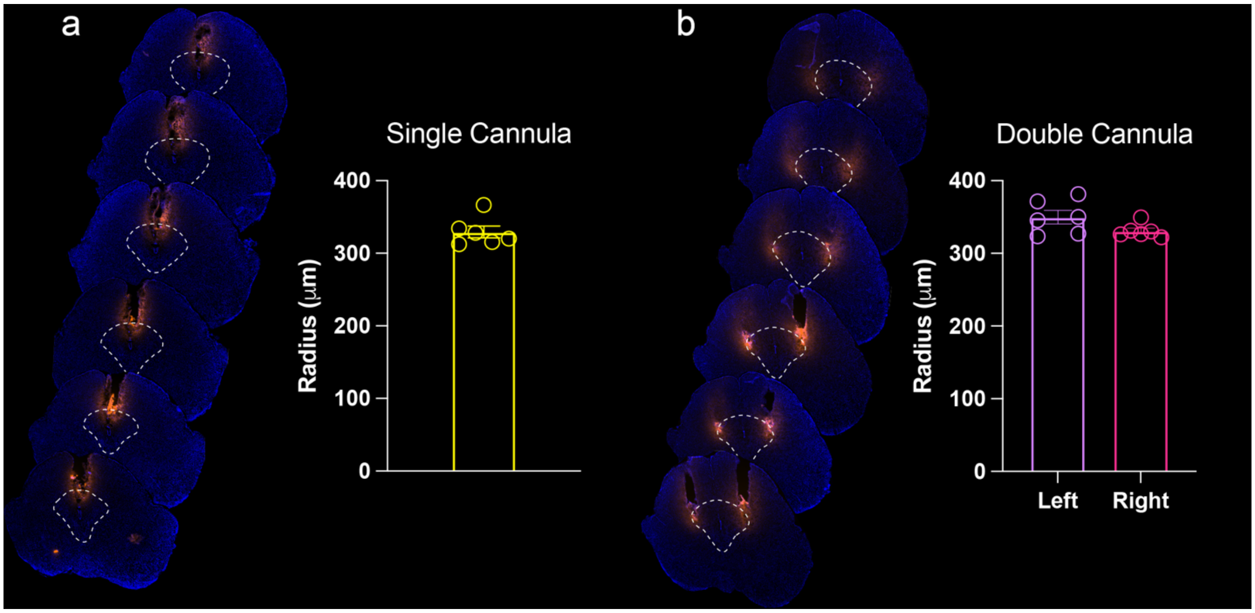


**Supplementary Fig. 15. PAG microinjection of fluorescently labeled muscimol for visualization of injection spread using the same injection coordinates, rate and volume as performed for the ligand microinfusions. a** Representative images for microinjection into the dmPAG (n = 6) showing minimal crossover to the lPAG. The average radius of spread was calculated to be 329.4 ± 19.83 µm. **b** Representative images for microinfusion into the lPAG (n = 6) showing minimal crossover into the dmPAG. The average radius of spread was calculated to be 349.8 ± 23.26 µm for the left hemisphere and 330.8 ± 9.69 µm for the right hemisphere. Post-experiment infusion with fluorescent muscimol allowed us to visualize cannula placements that were within the intended target region. Mice with cannula placements outside of the dmPAG or lPAG in the respective cohort were excluded. Data are shown as mean ± SEM.

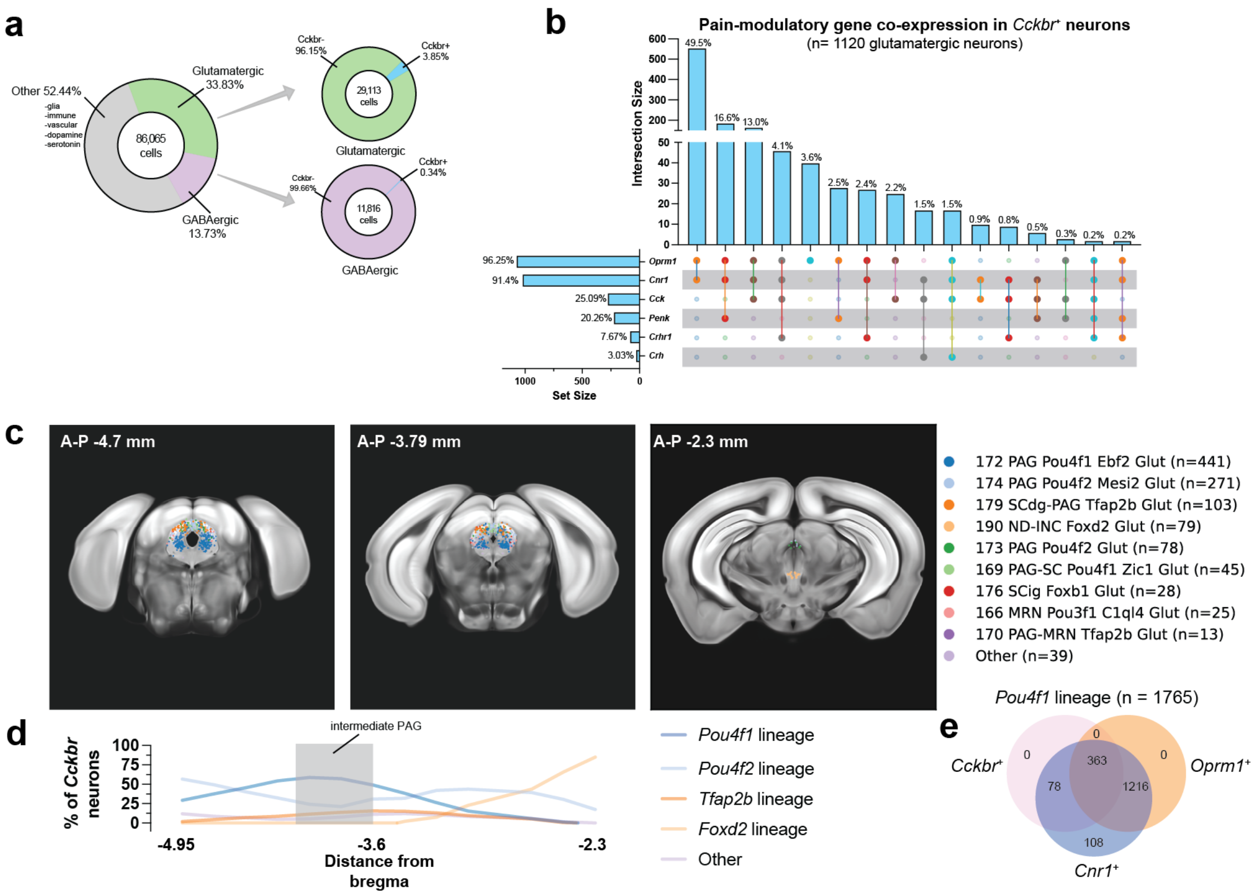


**Supplementary Fig. 16. Molecular identity and anatomical distribution of *Cckbr* neurons in the periaqueductal gray.** Single cell transcriptomic data from the Allen Brain Cell Atlas were used to map *Cckbr* expressing neurons within the PAG and characterize their molecular phenotype. **a** Overall cellular composition of the PAG and relative abundance of *Cckbr* expression. *Cckbr* was rare across all PAG cells and confined largely to neurons. When restricted to neuronal populations, expression was preferentially observed in glutamatergic neurons (n = 1120), compared to GABAergic neurons (n = 40). **b** Intersection analysis revealed that glutamatergic *Cckbr* neurons are overwhelmingly characterized by co-expression of *Oprm1* and *Cnr1*, with nearly half of neurons expressing both receptors simultaneously. In contrast, neuropeptide and stress-related genes (*Cck*, *Penk*, *Crhr1*, *Crh*) were present in substantially smaller subsets, indicating that this population is molecularly defined primarily by convergent opioid and cannabinoid signaling rather than broad engagement of multiple modulatory pathways. **c** Spatial distribution of lineage-defined *Cckbr* neurons across the rostrocaudal PAG. *Cckbr* cells were concentrated in intermediate PAG levels and were dominated by the *Pou4f1* lineage, with smaller contributions from *Pou4f2*, *Tfap2b*, and *Foxd2* lineages. The clustering of *Pou4f1*-derived neurons within the dorsolateral and lateral PAG identifies a discrete excitatory PAG population positioned to influence descending pain modulation. **d** Quantification of lineage composition along the rostrocaudal axis shows enrichment of the *Pou4f1* lineage in the intermediate PAG relative to other molecular subclasses. **e** A substantial fraction of the *Pou4f1* lineage population (n = 1765) co expresses opioid and cannabinoid receptors, demonstrating that *Cckbr* neurons occupy a molecular intersection of major endogenous analgesic systems.

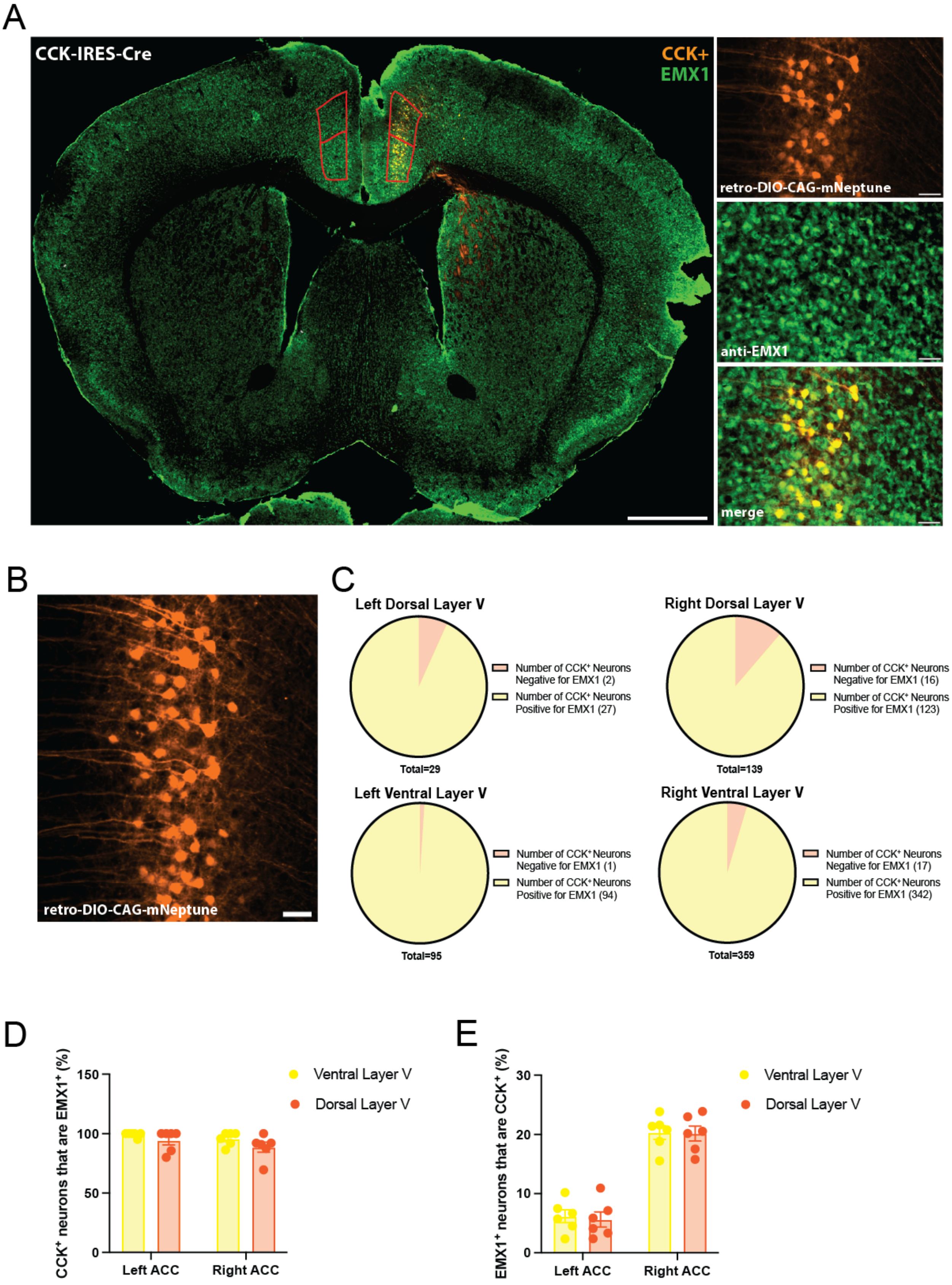
A
CCK-IRES-Cre
CCK+
EMX1
retro-DIO-CAG-mNeptune
anti-EMX1
merge
B
retro-DIO-CAG-mNeptune
C
Left Dorsal Layer V
Number of CCK+ Neurons Negative for EMX1 (2)
Number of CCK+ Neurons Positive for EMX1 (27)
Total=29
Right Dorsal Layer V
Number of CCK+ Neurons Negative for EMX1 (16)
Number of CCK+ Neurons Positive for EMX1 (123)
Total=139
Left Ventral Layer V
Number of CCK+ Neurons Negative for EMX1 (1)
Number of CCK+ Neurons Positive for EMX1 (94)
Total=95
Right Ventral Layer V
Number of CCK+ Neurons Negative for EMX1 (17)
Number of CCK+ Neurons Positive for EMX1 (342)
Total=359
D
CCK+ neurons that are EMX1+ (%)
150
100
50
0
Left ACC
Right ACC
Ventral Layer V
Dorsal Layer V
E
EMX1+ neurons that are CCK+ (%)
30
20
10
0
Left ACC
Right ACC
Ventral Layer V
Dorsal Layer V

**Supplementary Fig. 17. Co-expression of mNeptune and EMX1 in anterior cingulate cortex (ACC) cells following retrograde viral injection in the lPAG of CCK-Cre mice. a** Micrographs illustrate tissue from CCK-IRES-Cre mice that were retrogradely labled with viral tracer rAAV-CAG-DIO-mNeptune (orange, n = 3) and immunostained using an antibody against EMX1 (green). *Large panel:* A low magnification whole brain section highlights the specificity of CCK projections, showing axons traversing the internal capsule. Scale bar indicates 1 mm. *Smaller panels:* $CCK^+$ neurons robustly co-express EMX1 (yellow in merged image) in layer V of the ACC. Scale bar indicates 50μm. **b** Micrograph shows the pyramidal morphology of the retrogradely labeled cells. Scale bar indicates 50μm. **c** Circle charts show the total number of $CCK^+$ neurons in the left and right ACC for the dorsal and ventral subregions of layer V. The total number of $CCK^+$ neurons were then subdivided into those that negative for EMX1 or positive for EMX1. The retrograde viral injection was performed on the right lPAG, hence the greater number of CCK+ neurons on the right side. **d** The percentage of $CCK^+$ neurons that co-expressed EMX1 is presented for the ventral and dorsal subregions of layer V of the anterior cingulate cortex (ACC) in both hemispheres. **e** The percentage of $EMX1^+$ cells that co-expressed the viral tracer is presented for the ventral and dorsal subregions of layer V of the anterior cingulate cortex (ACC) in both hemispheres. Data are shown as mean ± SEM.

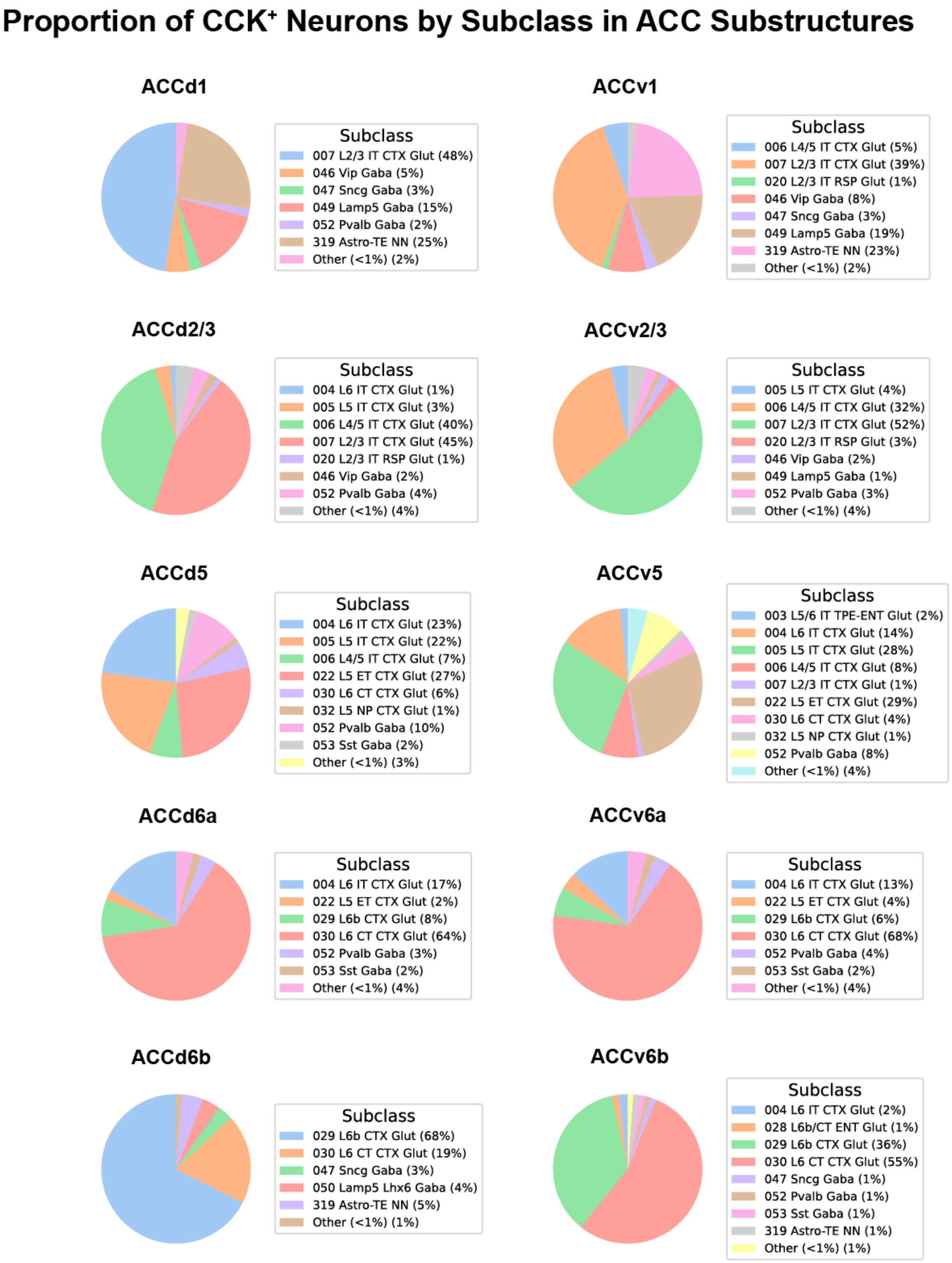


**Supplementary Fig. 18. Distribution of CCK-expressing neurons across anterior cingulate cortex (ACC) subclasses.** Proportion of Cck-expressing neurons across individual anterior cingulate cortex subregions (ACCd1, ACCv1, ACCd2/3, ACCv2/3, ACCd5, ACCv5, ACCd6a, ACCv6a, ACCd6b, ACCv6b), derived from the Allen Brain Cell Atlas spatial transcriptomic dataset. Each pie chart shows the distribution of transcriptomic neuron subclasses within each region, including excitatory intratelencephalic-like (IT) and extratelencephalic-like (ET) projection populations together with GABAergic interneurons. The "other" category groups subclasses that each accounted for less than 1% of *Cck*-expressing cells. Across subregions, *Cck* expression was dominated by excitatory projection neuron populations, with prominent representation of layer 2/3 and layer 5 IT-like classes and region-dependent contributions from deeper-layer projection types, whereas inhibitory subclasses formed a minor fraction. Differences between dorsal and ventral divisions indicate regional specialization of cortical CCK output populations. Percentages indicate the fraction of *Cck*$^{+}$ cells within each subclass.

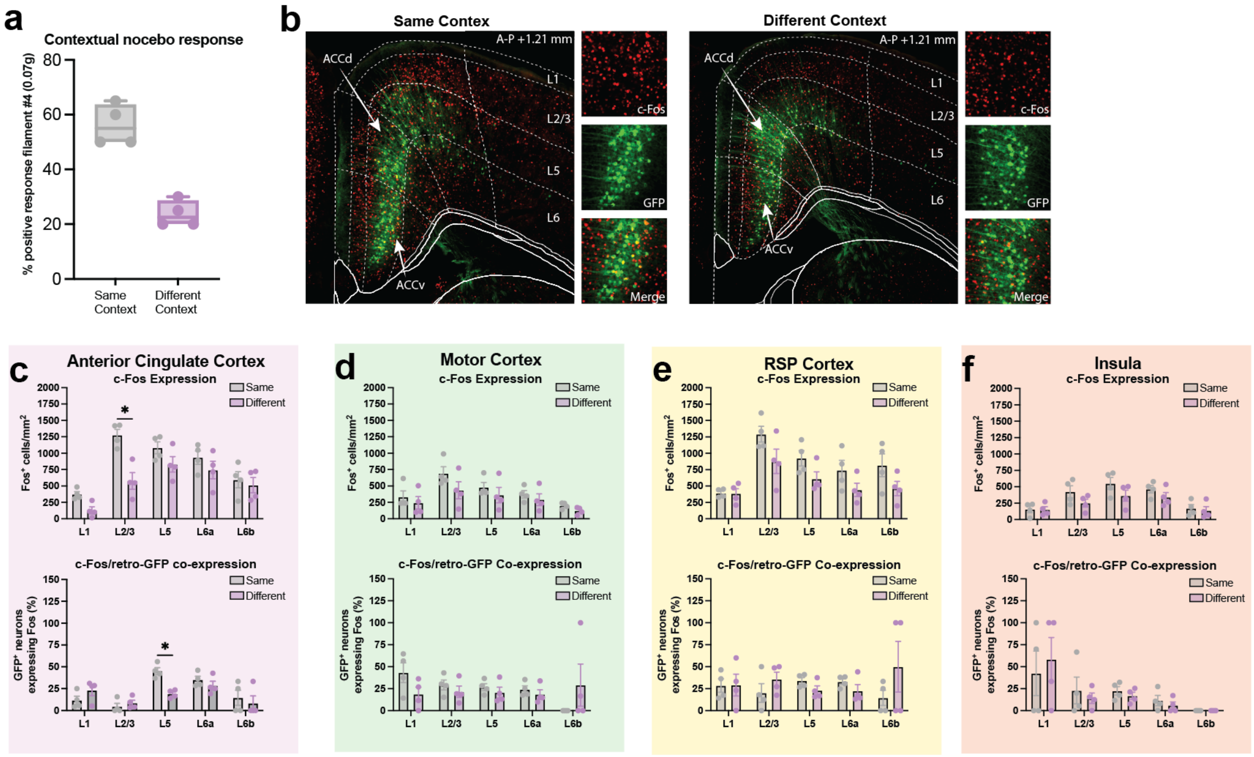


**Supplementary Fig. 19. Cortical activation during contextual nocebo expression. a** Mice tested in the same context (n = 4) show a higher percentage of withdrawals responses to a 0.07g von Frey filament than mice tested in a different context (n = 4) (Welch's t-test; t(5.09) = 7.3, p<0.001). **b** Representative images from the anterior cingulate cortex (ACC; A-P +1.21 mm) showing c-Fos immunoreactivity (red) and retrograde PAG-projecting neurons labeled with GFP (green) following testing in the same or a different context. Insets show higher-magnification view and merged signal. **c**–**f** Quantification of c-Fos expression (top) and c-Fos/retro-GFP co-expression (bottom) across cortical regions and layers: anterior cingulate cortex (**c**), motor cortex (**d**), retrosplenial cortex (**e**), and insular cortex (**f**). Mice tested in the same context showed increased overall neuronal activation in the ACC, driven primarily by superficial layer 2/3 neurons, whereas, co-expression analysis revealed selective recruitment of PAG-projecting ACC neurons within layer 5, indicating context-dependent engagement of the ACC→lPAG circuit rather than global cortical activation. Statistics include two-way ANOVAs, which revealed a significant interaction in the ACC for total c-Fos expression (context × layer interaction; F(1.92,11.54) = 9.205, p = 0.004) and c-Fos/retro-GFP expression (context × layer interaction; F(2.29,13.38) = 3.72, p=0.04), No interaction was detected in motor, retrosplenial, or insular cortex (all p's>0.05).
